# A genetic and microscopy toolkit for live imaging of whole-body regeneration in *Macrostomum lignano*

**DOI:** 10.1101/2024.04.28.590365

**Authors:** R. Nelson Hall, Hongquan Li, Chew Chai, Sidney Vermeulen, Robin R. Bigasin, Eun Sun Song, Jesse Gibson, Manu Prakash, Andrew Z. Fire, Bo Wang

## Abstract

Live imaging of regenerative processes can reveal how animals restore their bodies after injury through a cascade of dynamic cellular events. Here, we present a comprehensive toolkit for live imaging of whole-body regeneration in the flatworm *Macrostomum lignan*o, including a high throughput cloning pipeline, targeted cellular ablation, and advanced microscopy solutions. Using tissue-specific reporter expression, we examine how various structures regenerate. Enabled by a custom, low-cost luminescence/fluorescence microscope, we overcome intense stress-induced autofluorescence to demonstrate the first application of genetic cellular ablation in flatworms to reveal the limited regenerative capacity of neurons and their essential role during wound-healing. Finally, we build a novel open-source tracking microscope to continuously image moving animals throughout the week-long process of regeneration, quantifying kinetics of wound healing, nerve cord repair, body regeneration, growth, and behavioral recovery. Our findings suggest that nerve cord reconnection operates independently from primary body axis re-establishment and other downstream regenerative processes.

## Introduction

In response to injury, regenerative animals initiate a cascade of dynamic cellular processes, including signaling, apoptosis, proliferation, differentiation, and patterning to restore lost tissue (Gemberling et al., 2013; Tanaka, 2016; Reddien, 2018). However, most studies infer dynamics based on snapshots spaced out in time. In contrast, live imaging can allow for continuous direct observation of these dynamic processes in the same individual (Currie et al., 2016; Zattara et al., 2016; Ratnayake et al., 2021). While short-term live imaging is routine, imaging the entire process of regeneration has been challenging, if not impossible, for three major reasons. First, very few systems offer both optical transparency and minimal autofluorescence required for fluorescence imaging. Second, except for a few genetically tractable systems (Siebert et al., 2019; Ricci & Srivastava, 2021; Weissbourd et al., 2021; Paix et al., 2023), most regenerative organisms lack tissue-specific transgenic labeling necessary for visualizing cells *in vivo*. Finally, restraining, or paralyzing animals for prolonged imaging over days – the timescale relevant for observing regeneration – is technically challenging and can negatively affect the organism’s physiology and change the course of regeneration. In this study, we address these challenges by combining the favorable physiology and genetics of the flatworm *Macrostomum lignano* with novel microscopy techniques to enable sensitive and long-term imaging of whole-body regeneration and illustrate the mechanistic insights that may be gained through such studies.

The interstitial, marine flatworm *M. lignano*, studied for diverse biological processes (Wudarski et al., 2020) including sexual selection (Brand et al., 2020; Marie-Orleach et al., 2021), bio-adhesion (Lengerer et al., 2014), genome evolution (Wasik et al., 2015; Zadesenets et al., 2017; Wiberg et al., 2023), and host-microbiome interactions (Ma et al., 2023), is also capable of regenerating all tissues posterior to the pharynx but not anterior structures, presenting a unique opportunity to compare the molecular and cellular basis of regenerative and non-regenerative outcomes (Egger et al., 2006, 2009). In addition, *M. lignano* is conducive to live imaging thanks to its small body size, optical transparency, minimal autofluorescence, and robust physiology.

In contrast to other commonly studied flatworm models like planarians, which have limited transgenic tools (Hall et al., 2022), *M. lignano* has the ability to integrate exogenous DNAs into its genome, enabling transgenesis (Wudarski et al., 2017; Ustyantsev et al., 2021; Wudarski et al., 2022), which is further facilitated by the animal’s short egg-to-egg generation time (3 weeks at 20 ℃) and abundant, injectable zygotes (Mouton et al., 2018). However, our repertoire of transgenic tools is in its infancy and advanced microscopy tools capable of imaging entire organisms in their native physiological state throughout regeneration have been lacking. Here, we integrate three key technologies to synergistically facilitate the use of *M. lignano* as a model system for the study of regeneration biology.

First, we established and utilized a modular library of genetic parts to enable the hierarchical assembly of complex expression cassettes within a single day. This high throughput cloning is coupled with a refined injection procedure allowing a single person to comfortably inject ∼200 fertilized eggs within a 2 hour session, which has routinely provided us with 1-2 germline-transmitting transformants per session. Using this pipeline, we have generated tissue-specific reporter lines expressing green nanolantern (GeNL) (Suzuki et al., 2016) which allows for both fluorescence and luminescence imaging. These lines have allowed us to better characterize anatomical structures, track the dynamics of axonal extension during neural regeneration, and reveal key steps during the regeneration of complex organs such as the male copulatory apparatus.

Second, we demonstrate the first application of targeted ablation in a regenerative flatworm by generating tissue-specific lines expressing nitroreductase (NTR2.0) (Sharrock et al., 2022), a genetically encoded, inducible toxin, but found that intense stress-induced autofluorescence resulting from cellular ablation obscured the ablation outcomes. Therefore, we built an affordable multimodal epifluorescence/luminescence microscope that circumvented autofluorescence and enabled the sensitive detection of cells to reveal a lack of neural regeneration following ablation, indicating that the turnover of the *M. lignano* nervous system may be slow, if it occurs at all. Furthermore, neural-ablated animals failed to heal their wounds after amputation, suggesting that the nervous system plays a critical role in the organism’s ability to repair and regenerate.

Third, we designed and built an epifluorescence tracking microscope, based on the open-source Squid platform (Li et al., 2020), to image live, free-moving animals for up to one week. Using infrared (IR) light to track the animal, this microscope continually adjusts the stage position to ensure the subject remains within the field-of-view (FOV). With this, we demonstrated the first continuous fluorescence imaging of whole-body regeneration and identified milestones during posterior neural regeneration with high temporal and spatial resolution. A particularly striking observation is that the rate of nerve cord extension depends on the distance needed to travel for nerve cords to reconnect after asymmetric amputations.

Finally, combining all these techniques, we investigated the function of *β-catenin*, a central regulator of the Wnt signaling pathway, in *M. lignano* regeneration. We found that animals failed to extend their body and regenerate posterior structures upon *β-catenin* RNAi, but nerve cord reconnection was unaffected, suggesting that nerve cord repair is separated from other processes of regeneration, including the primary body-axis specification. With the toolkit we created, our work positions *M. lignano* to become a powerful platform for live imaging studies to reveal dynamic cellular processes during tissue regeneration.

## Results

### A modular cloning protocol enables rapid generation of tissue-specific reporter lines

Cloning can often be a bottleneck when constructing complex transgene cassettes. To streamline this process and facilitate resource sharing, we took inspiration from the idea of modular assembly, which breaks down transgenes into ‘parts’ – promoters, genes, and terminators – that can be hierarchically assembled through simple biochemical reactions (Canton et al., 2008). Building upon the 3G Assembly pipeline (Halleran et al., 2018), we have implemented a single-day protocol to generate multi-transcriptional unit (TU) plasmids. We first assembled parts via Golden Gate into TUs, which were further assembled into plasmids via Gibson assembly using positionally indexed homology arms added during the prior Golden Gate reactions (**Figure 1A**). Our approach minimizes scar sites within TUs but preserves modularity and flexibility. A key change we made to the original 3G method is the introduction of a new backbone, into which all parts can be cloned, thereby reducing the reliance on PCR for generating part fragments. This allows for the long-term storage of parts as glycerol stocks and simplifies the process of part sharing and reuse. We have built a parts library (**Table S1, File S1**) and expect to continue expanding it.

**Figure 1:**
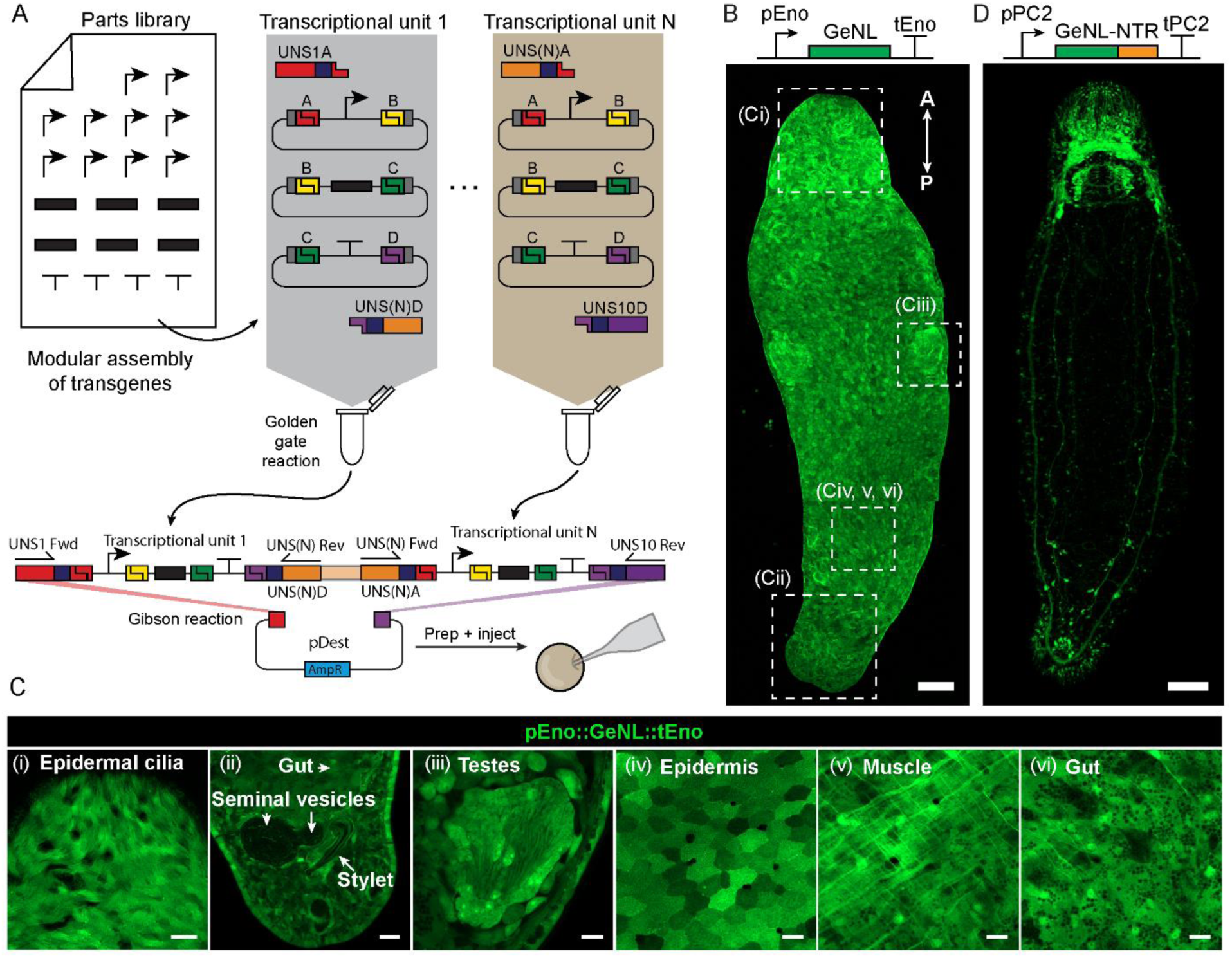
A modular cloning protocol enables rapid generation of tissue-specific reporter lines. **(A)** A diagram showing the design of the modular cloning toolkit. A parts library, composed of promoters, genes, and terminators are assembled into different TUs via Golden Gate reactions (gray and brown regions), along with adapter oligos containing numbered unique sequences (UNS) which define their position in the later Gibson assembly. Within each reaction, each lettered site (A through D) ligates with its complementary overhang (matching colors), thus stitching together a complete TU flanked by the appropriate unique sequence adapters. A PCR reaction amplifies the assembled TU and Gibson assembly then combines TUs into a destination vector (pDest). Throughout this paper, transgenes are presented in the form ‘pPromoter::Gene::tTerminator’. **(B)** Confocal image showing mNeonGreen fluorescence in the pEnolase::GeNL::tEnolase strain. Dashed boxes correspond to the anatomical regions depicted in panel C. A-P: anterior-posterior. Scale bar: 100 µm. **(C)** Example images of the pEnolase::GeNL::tEnolase transgene expression across various tissues: (i) epidermal cilia; (ii) tail, showing gut, seminal vesicles, and stylet; (iii) testes; (iv) epidermis; (v) muscle; (vi) gut. Images are single confocal slices. Scale bars: 20 µm (i, ii), 10 µm (iii-vi). **(D)** Confocal image of an animal expressing pPC2::GeNL-NTR2.0::tPC2. Scale bar: 100 µm.

Using our toolkit, we have created a series of transgenic lines carrying increasingly complex expression cassettes. First, we generated lines ubiquitously expressing GeNL, a fusion of mNeonGreen and nanoluciferase (Nluc) (Suzuki et al., 2016), under the control of either the enolase (**Figure 1B**) or the eukaryotic elongation factor 1α (Eef1α) promoter. Notably, the pEnolase::GeNL::tEnolase line exhibited more uniform expression, at least in the epidermis (**Figure 1C**, **Figure S1A**), allowing us to visualize various anatomical features including the epidermal cilia, seminal vesicles, stylet, testes, muscles, and intestinal cells (**Figure 1C**, **Video S1**).

Next, we created a transgenic line co-expressing GeNL and NTR2.0 in neurons, driven by the promoter of a prohormone convertase 2 (PC2) homolog (**Figure 1D**) – a gene expressed across all neural cell types in *M. lignano* (**Figure S1B).** The pPC2::GeNL-P2A-NTR2.0::tPC2 line showed strong transgene expression in the central nervous system, consisting of the cephalic ganglia, two major and various minor ventral nerve cords, as well as peripheral neural structures such as the anterior sensory organs and posterior neurons associated with reproductive and adhesive organs. The NTR gene was incorporated to enable targeted chemical ablation of the labeled cells, which is discussed below.

Finally, we produced lines with transgenes expressed in multiple distinct patterns. We used the APOB and MHY6 promoters (Wudarski et al., 2017) to co-express GeNL and NTR2.0 in intestinal and muscle cells, respectively (**Figure S2A-B**). The constructs also contained an additional expression cassette, pEef1α::mScarlet::tEef1α, serving as a co-transfection marker. This marker was crucial in determining the locations of GeNL^+^ cells in various organs (as described below). Overall, these results demonstrate how our approach can combine new and existing genetic parts and facilitate the efficient generation of genetic lines carrying complex transgenes.

#### Tissue-specific labeling resolves detailed anatomy

These tissue-specific reporter lines allowed us to characterize the fine anatomy of various tissues in live animals using confocal microscopy. In the PC2 reporter line, the cephalic ganglia was clearly labeled in the anterior, revealing a dense neuropil located just above the photoreceptors, and a battery of ciliated sensory neurons projecting towards the far anterior of the animal (**Figure 2A**). The pharynx was encircled by a highly enervated ring (**Figure 2A**, **Video S2**), and two photoreceptors each projected an axon into the neuropil (**Figure 2B**). Along the body, we observed a pair of major ventral nerve cords and several minor nerve cords situated medially, which often projected horizontally to connect between nerve cords (**Figure 2C**). The tail region was highly innervated, with the ventral nerve cords looping around the tail base, encircling a mesh of neurons in the middle. Along the exterior of the tail was a fan-like array of neurons likely controlling the duo-adhesive gland of the adhesive organ (**Figure 2D**). A closer inspection of these neurons revealed that they were connected to the major nerve cord by a single long projection with a second projection extending to the edge of the tail (**Figure 2E**). Finally, we discovered unique arrangements of neurons around the stylet opening (**Figure 2F**) and the antrum (**Figure 2G**), the opening from which eggs are deposited. Intriguingly, similar groupings of neurons regulate various copulatory behaviors and egg laying from the vulva in *C. elegans* as well (Emmons, 2018). Notably, many of these neural structures were not visible in previous studies of the nervous system using immunofluorescence (Ladurner et al., 1997; Morris et al., 2007).

**Figure 2:**
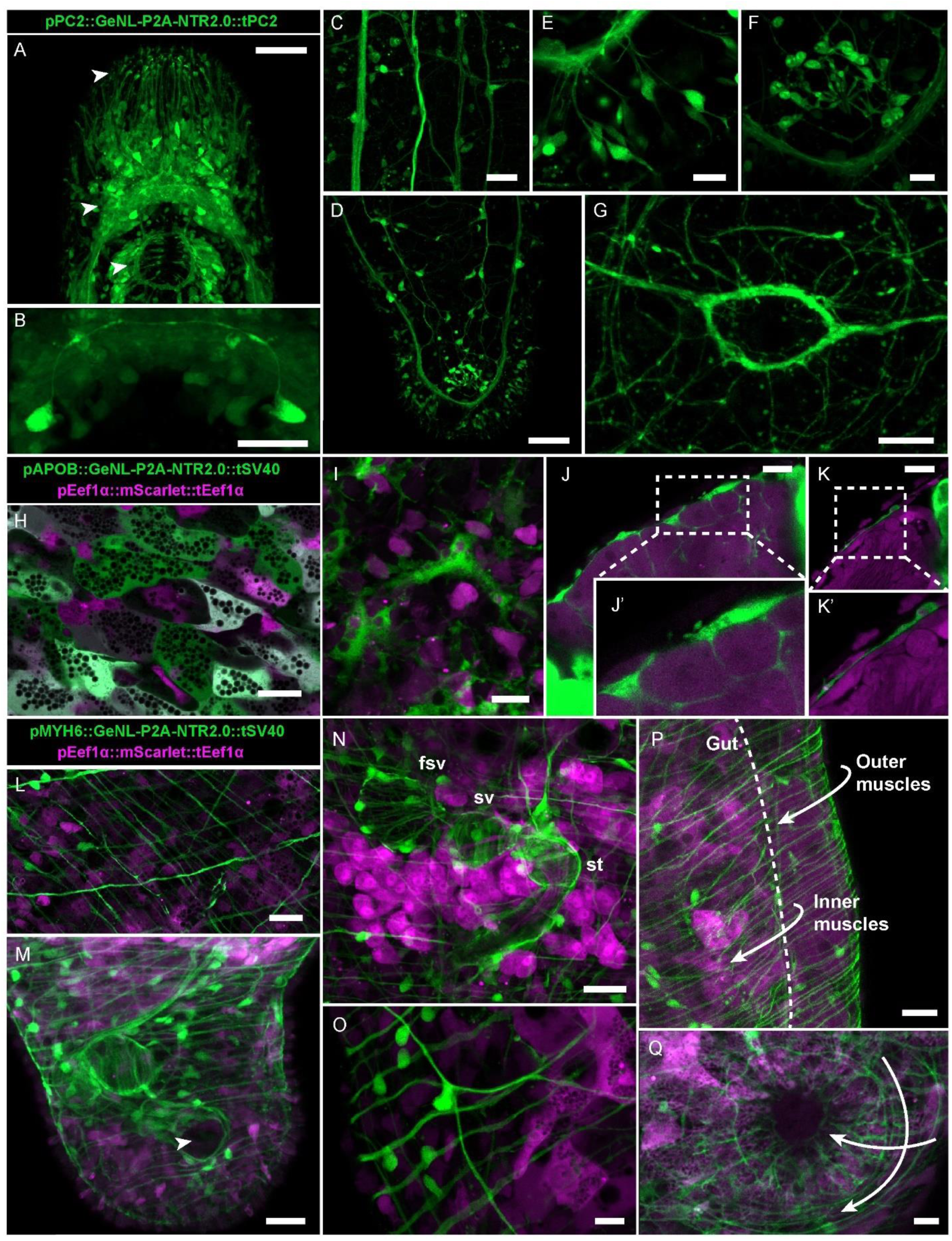
Tissue-specific labeling resolves detailed anatomy. **(A)** Confocal image of the *M. lignano* brain. Arrows highlight (from top to bottom) anterior sensory projections, the dense neuropil, and pharynx. The imaging regions of all panels in this figure are specified in **Figure S1C**. **(B)** A close-up of the two photoreceptor cells and the axonal projections they send into the neuropil. **(C)** The major nerve cord (left) and minor nerve cords (right) run from the head to the tail of the animal. **(D)** An overview of the nervous system in the tail showing the major nerve cords looping around the base of the adhesive organ. **(E)** Neurons of the adhesive organ each sending a projection into the major nerve cord and another outward to the epidermis. **(F)** A circular arrangement of neurons around the opening of the stylet. **(G)** Dense nerve fibers around the opening of the antrum. **(H)** Tiled gut cells with highly vacuolated cytoplasm. **(I)** Large pAPOB::GeNL^+^ cells (green) enclosing other cells (magenta) present in the anterior of the animal. **(J)** Ovaries (magenta) are wrapped by pAPOB::GeNL^+^ cell bodies (green) on the exterior, extending cytoplasmic processes around individual oocytes. Inset: two oocytes surrounded by cytoplasmic processes of pAPOB::GeNL^+^ cells. **(K)** A pAPOB::GeNL^+^ cell (green) sits on the exterior of the testes (magenta). Inset: a magnified view of this cell, which extends a long process down the length of the testes. **(L)** Crosshatched circular and diagonal muscle cells (green) with intestinal cells beneath (magenta). **(M)** An overview of the tail, showing the muscular structure of the male copulatory apparatus (green) within. Arrow: the stylet opening. **(N)** The male copulatory apparatus, with the false seminal vesicle (fsv), seminal vesicle (sv), and stylet (st) visualized by the layer of muscle cells (green) wrapping each compartment. Abundant prostate gland cells (magenta) surround the seminal vesicles and stylet. **(O)** Individual muscle cell bodies hang like beads from circular muscle fibers (green) around the gut (magenta). **(P)** Two layers of muscles with body-wall muscles under the epidermis and another around the gut. The ovaries (magenta) sit between these two muscle layers (green). Dashed line: boundary between the gut and parenchyma. Arrows trace the two layers of muscles. **(Q)** The antrum is surrounded by concentric rings of muscle intersected by a second set of perpendicular radial muscle cells (green). Arrows trace the radial and perpendicular muscle fibers. Scale bars: 50 µm (**A**, **D**), 20 µm (**B**, **C**, **G**, **H**, **L**-**O**, **Q**), 10 µm (**E**, **F**, **I**-**K**, **P**).

In the APOB reporter line, the gut architecture was revealed as a dense tiling of highly vacuolated cells. While both APOB and Eef1α promoters drove expression in the gut, the ratio of their expression was variable between cells (**Figure 2H**), consistent with our single-cell ATAC-seq data suggesting that *eef1α* is highly expressed in one subpopulation of intestinal cells (**Figure S1B**). In the head, large pAPOB::GeNL^+^ cells extended numerous tracts of cytoplasmic processes, which often encased neighboring cells (**Figure 2I**). Surprisingly, pAPOB::GeNL^+^ cells were present within the ovaries, with their cell bodies located at the periphery and cytoplasmic processes surrounding developing oocytes (**Figure 2J, 2J’**). Similarly, we observed pAPOB::GeNL^+^ cells straddling the exterior of the testes, though not within the testes (**Figure 2K, 2K’**). These observations are consistent with single-cell analysis that revealed a population of *apob*^+^ *cathepsin*^+^ cells (**Figure S1B**), suggesting that these previously undescribed extraintestinal cells could play phagocytic roles patrolling through various organs.

The MHY6 reporter line showed distinct layers of circular and diagonal muscle fibers beneath the epidermis (**Figure 2L**) (Rieger et al., 1994). The tail was particularly rich in muscle fibers (**Figure 2M**). A closer examination revealed that the seminal vesicle, false seminal vesicle, and stylet were all wrapped in muscular rings, which likely control sperm expulsion and stylet movement during copulation (**Figure 2N**). In parallel, the pEef1α::mScarlet expression uncovered clusters of presumptive prostate gland cells (Ladurner et al., 2005), extending into the stylet (**Figure 2N**, **Figure S2C**). Within the trunk, a secondary inner layer of muscle fibers pressed against internal organs including the gut (**Figure 2O**). This was most obvious around the testes and ovaries, which were sandwiched between the muscle layers (**Figure 2P**, **Video S3**). Finally, the antrum was surrounded by concentric muscle rings (**Figure 2Q**), aligning with the neural ring (**Figure 2G**), suggesting that the muscles are innervated to regulate egg laying. Together, these observations demonstrate the capability to resolve anatomical details with high resolution using combinations of tissue-specific and ubiquitous transgenes, establishing many anatomical landmarks that should be instrumental in evaluating the regeneration process.

### Time course imaging using tissue-specific reporters reveals stages of posterior regeneration

To investigate how tissue structures regenerate, we amputated each transgenic line (PC2, APOB, and MYH6) and performed live imaging at regular time points over a week. Using the PC2 line, we observed that the major nerve cords remained severed and produced an elaborate fan-like array of axonal projections towards the wound at 6 hours post-amputation (hpa) (**Figure 3A**). Occasionally, these major nerve cords connected with minor ones forming loops (**Figure 3B**). By 48 hpa, the nerve cords were fully reconnected, though no adhesive gland neurons were yet observed (**Figure 3C**). By 4 days post-amputation (dpa), we observed an increasingly elaborate array of neurons at the tail tip (**Figure 3D**), consistent with prior observations that the duo-adhesive system regenerates by 3 dpa (Egger et al., 2006). By 7 dpa, the tail plate regained its normal club-shaped appearance (**Figure 3E**). These results suggest a potential role for axonal guidance cues during wound healing as existing nerve cords appear to seek out for connection, followed by neurogenesis in the tail plate as the adhesive system regrows.

**Figure 3:**
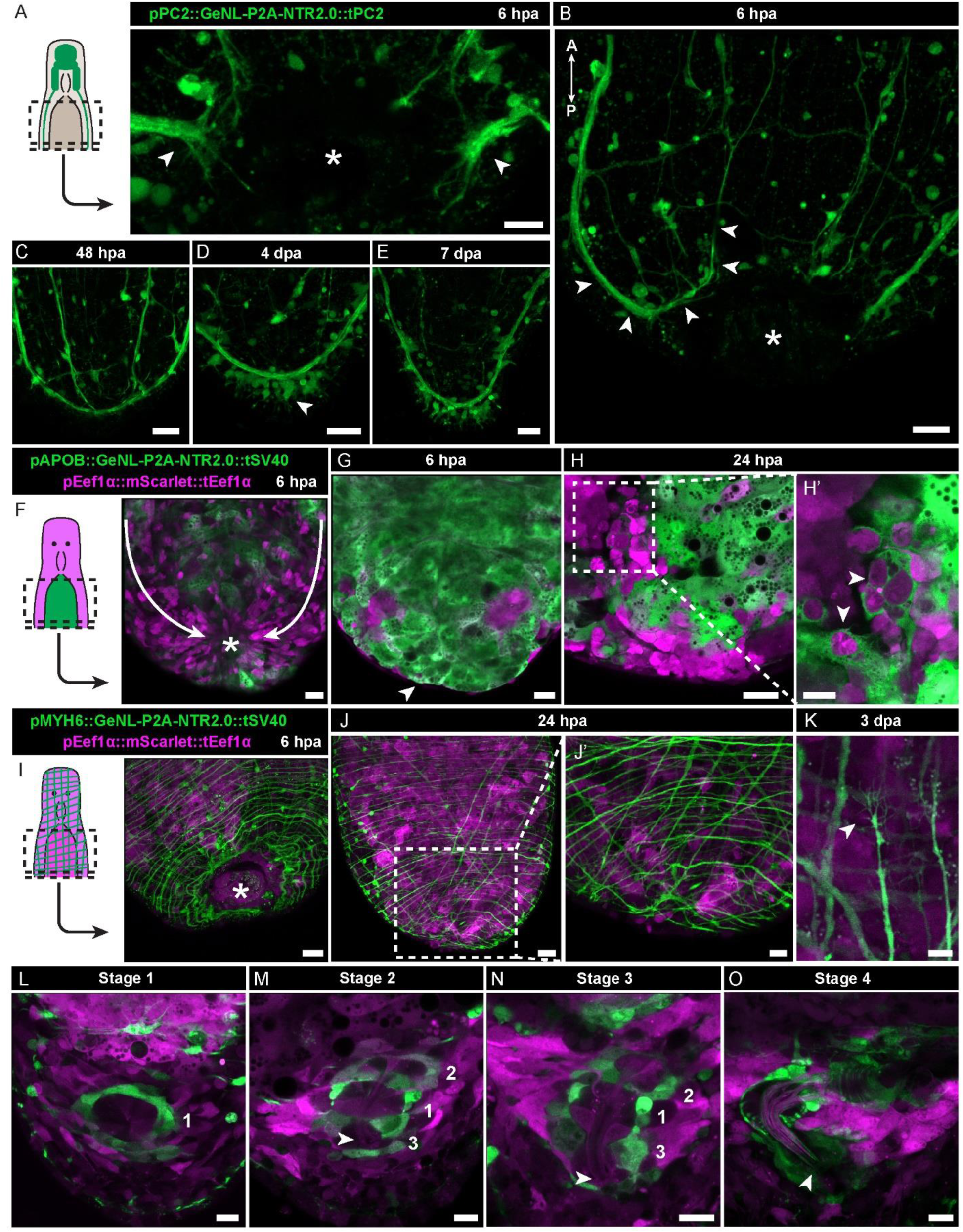
Time course imaging using tissue-specific reporters reveals stages of posterior regeneration. **(A)** Cartoon showing an amputated animal with the nervous system highlighted (left). The images are taken from the head fragment. At 6 hpa, the major nerve cords (arrows) are severed but extend projections towards the wound site (asterisk). Dashed box: region imaged. **(B)** Another view of severed ventral nerve cords extending projections towards the wound (asterisk) forming loops with minor ventral nerve cords (arrows trace the loop). **(C)** By 48 hpa, the nerve cords have reconnected fully. **(D)** By 4 dpa, neurons of the adhesive organ have begun repopulating the tail. **(E)** By 7 dpa, the adhesive organ has regained its normal shape. **(F)** Cartoon showing an amputated animal with GeNL expression in the gut (green) and ubiquitous mScarlet expression (magenta) (left). By 6 hpa, the epithelium (magenta) is stretched towards the wound with the gut underneath (green). Dashed box: region imaged. **(G)** A confocal slice deeper into the tissue showing the gut (green) appears immediately beneath the epithelium (magenta) at the wound. **(H)** By 24 hpa, phagocytes adjacent to the gut (green) are in close contact with cells in the regenerating blastema (magenta) (left). Inset: a magnified view showing numerous blastemal cells (magenta) are surrounded by cytoplasmic processes of pAPOB::GeNL^+^ cells (green) (right). **(I)** Cartoon showing an amputated animal with GeNL expression in muscles (green) and ubiquitous mScarlet expression (magenta) throughout the body (magenta) (left). At 6 hpa, circular muscles are wavy and buckled as the wound (asterisk) closes. Dashed box: region imaged. **(J)** By 24 hpa, the muscles (green) enclose the wound (left). Inset: a magnified view showing slight disorganization in the new meshwork of muscle fibers (green) in the blastema (right). **(K)** An example of a muscle fiber at 3 dpa with a terminus ending in many filamentous projections (green). (**L-O**) A staged progression of male reproductive organ regeneration. **L**, stage 1, a ring of GeNL^+^ cells appears in the posterior blastema. **M**, stage 2, multiple rings of GeNL^+^ cells outline the primordia of seminal vesicles and stylet. A stylet primordium begins to form (arrow). **N**, stage 3, the rings continue to grow into larger chambers as the stylet continues to elongate and the prostate gland cells (magenta) extend projections into the chambers and growing stylet (arrow). **O**, stage 4, the stylet adopts its final bent shape, the chambers have grown into matured seminal vesicles, and numerous prostate gland cells send abundant processes into the stylet. Numbers: GeNL^+^ circular rings. Scale bars: 20 µm (**B-J**), 10 µm (**A**, **L-M, H’, J’**), 5 µm (**K**).

Turning to the APOB line, at 6 hpa, the epidermis at the wound appeared distorted and stretched, resembling a drawstring being pulled tight (**Figure 3F**). Below the wound, the gut sat directly beneath the epidermis with no tissue in between (**Figure 3G**). By 24 hpa, the blastema had grown between gut and epidermis, often with a subset of cells encircled by pAPOB::GeNL^+^ cells **(Figure 3H, 3H’**). This pattern was similarly observed for the pAPOB::GeNL^+^ cells within the head during homeostasis in the APOB line, hinting at a possible role of phagocytes in removing extra cells.

Finally, we analyzed the musculature during regeneration using the MYH6 line. At 6 hpa, muscle fibers formed wavy concentric rings around the wound (**Figure 3I**), and by 24 hpa, muscles enclosed the wound site (**Figure 3J, 3J’**). We often observed muscle fibers with termini forming multiple filamentous projections in both pre-existing and regenerating tissues, suggesting that the muscle network was in the process of reforming during regeneration (**Figure 3K**).

The MYH6 line also revealed key steps during the organogenesis of the male copulatory apparatus. Starting at 3 dpa, a ring of pMYH6::GeNL^+^ cells appeared in tail blastema with a small rosette of pEef1α::mScarlet^+^ cells at the center (**Figure 3L**). Next, this ring developed into three connecting rings that went on to become seminal vesicles and the stylet sheath, inside of which a small stylet primordium began to form (**Figure 3M**). The rings continued to grow, as did the stylet. By this point, many prostate gland cells were present around the edges of the rings and extended long processes down into the stylet, sometimes traveling many cell bodies in distance (**Figure 3N**, **Figure S2D**). Finally, the rings became fully formed chambers sheathed in muscle fibers, and the stylet adopted a sharp bend at its base that was heavily invaded by processes of the surrounding prostate glands (**Figure 3O**). These results highlight the complex dynamic processes within regenerating tissues, providing multiple examples of intricate cellular processes, which likely involve continuous coordination between cells and across tissues.

### Luminescence imaging tracks neural ablation outcomes

Cell type-specific expression not only facilitates the observation of tissue dynamics during regeneration but also enables targeted cell ablation using conditional, genetically encoded toxins like NTR2.0 (Sharrock et al., 2022), which can help elucidate the function of specific cell populations; however, such a method had never been applied in flatworms. To test the functionality of NTR2.0 in flatworms, we treated each of our strains with 0.5 or 5 mM metronidazole (MTZ), a prodrug reduced by NTR to induce cytotoxicity (Sharrock et al., 2022), for 7 days. Compared to controls treated with an equal concentration of DMSO (vehicle), the PC2 strain showed a loss of muscle tone and paralysis (**Video S4**), the APOB animals ejected their gut (**Figure S3A)**, and the MYH6 line, though still capable of moving, contracted into spherical shapes (**Video S4**). We chose the PC2 strain for further analysis, as they preserved their gross morphology after the MTZ treatment. However, we also observed drastically increased autofluorescence after treatment, obscuring the extent of neural ablation (**Figure 4A**).

**Figure 4:**
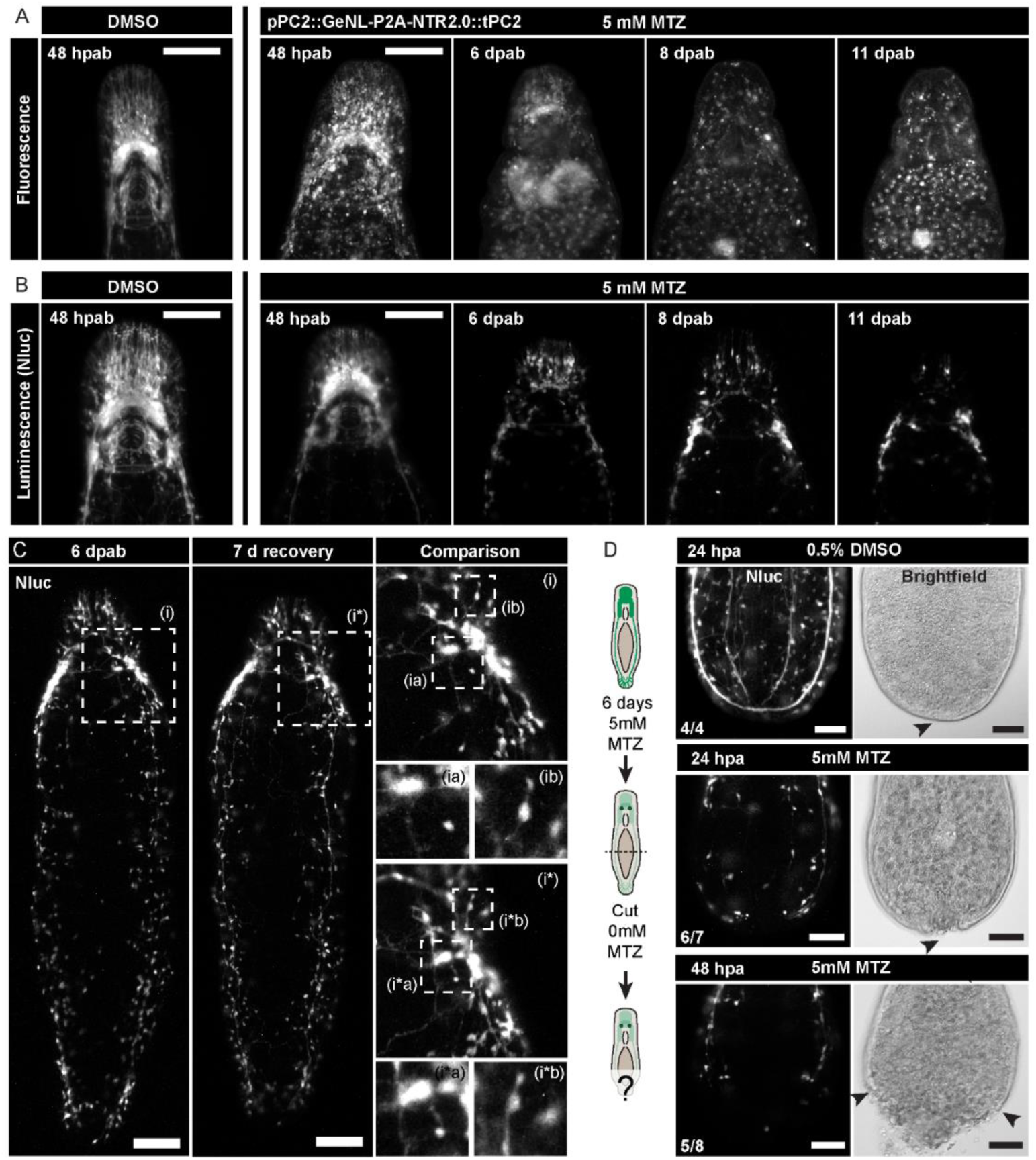
Luminescence imaging tracks neural ablation outcomes. **(A-B)** Time-course fluorescence (A) and luminescence (B) images of PC2 animals after neural ablation. **(C)** Images on the same animal immediately after 6 d MTZ (5mM) treatment (left) and after 7 days of recovery in ASW (middle). The highlighted regions (dashed boxes, i, i*) show little change between the two time points. Further magnified views (dashed boxes, ia, ib, i*a, i*b) show specific cell arrangements that can be mapped between the time points to highlight the lack of change (right). **(D)** Neural ablation prevents wound-healing and subsequent regeneration. Control animals treated with 0.5% DMSO successfully reconnected their nerve cords and healed the epidermis (top) (n = 4/4). Animals after neural ablation showed no ventral nerve cords and failed to heal the epidermis by 24 hpa (middle) (n = 6/7). By 48 hpa, epidermal integrity continued to deteriorate, resulting in animals with exposed wounds (bottom) (n=5/8). All ablated and amputated animals (n > 40) lysed by 7 dpa. Arrows: a normally closed epidermis (top, disrupted epidermis (middle), severely disrupted epidermis (bottom). Scale bars: 100 µm (**A-C**), 50 µm (**D**).

To overcome this challenge, we utilized the Squid platform (Li et al., 2020) to develop an upgraded version of our previous low-cost luminescence microscope (Hall et al., 2022) by incorporating full motorization, dual-color fluorescence imaging, and a python-based graphical user interface (**Figure S3B, Figure S4A)**. Instead of an EMCCD, this new microscope uses a cooled CMOS camera, with high peak quantum efficiency (91%) and very low dark current, reducing the camera cost by over an order of magnitude with little compromise in performance, and making luminescence imaging more broadly accessible. Comparing fluorescence and luminescence images of the same animal, we confirmed that luminescence imaging was consistent with fluorescence but had reduced background signal, permitting the visualization of fine neural processes (**Figure S3C)**.

In contrast to fluorescence, luminescence imaging revealed a progressive reduction of neuronal cells in the PC2 animals kept in 5mM MTZ (**Figure 4B**). By 11 days post-ablation (dpab), only a few neurons remained, yet our microscope was able to detect their faint luminescent signal. The ability to sensitively detect cells without confounding autofluorescence opens the possibility to explore the regenerative capacity of the nervous system post-ablation.

To investigate whether the nervous system could regenerate after ablation, we removed the animals from MTZ at 6 dpab, followed by 7 days of recovery in artificial seawater (ASW). We performed luminescence live imaging before and after the recovery and found no substantial changes to the degenerated nervous system, with landmark cells persisting across the whole period (**Figure 4C**). This indicates a lack of neural regeneration after extensive neural ablation.

Finally, we explored whether the nervous system is necessary for regeneration and whether injury is needed to activate neural regeneration after ablation. We amputated animals at 7 dpab and followed their regeneration in the absence of MTZ. MTZ-treated non-transgenic controls showed full regeneration by 7 dpa (**Figure S3D**). The PC2 strain treated with DMSO (vehicle) exhibited normal wound healing and the nerve cords reconnected by 24 hpa. In contrast, the PC2 animals subjected to neural ablation with MTZ failed to reconnect what remained of their nerve cords and the posterior wound remained open (**Figure 4D**), eventually leading to lysis. This observation highlights the critical role of the nervous system in wound healing and shows that injury alone does not trigger neural regeneration. Overall, these results demonstrate the first application of cellular ablation in regenerative flatworms and showcase the importance of luminescence imaging in accurately assessing ablation outcomes.

### Tracking microscope enables continuous imaging of posterior neural regeneration in free-moving animals

To observe regeneration from start to finish in real-time requires long-term imaging performed on minimally perturbed, freely moving animals. Using the Squid platform (Li et al., 2020), we built a novel epifluorescence microscope capable of tracking the animal while continuously capturing high resolution images.

The animal is imaged in bright field with low intensity infrared (IR) light, which does not elicit photophobic responses in *M. lignano* (Paskin et al., 2014) or heat up the water. The images are acquired at a frequency of ∼6 Hz and segmented in real-time to determine the animal’s centroid and adjust the stage’s position to keep the animal at the center of the FOV. In parallel, the microscope acquires fluorescence images through a separate optical path in up to four possible channels (**Figure 5A, Figure S4B**). To avoid the loss of tracking due to vertical movements of the animal, we engineered a flat imaging chamber that confined the animal in only the z-axis (**Figure 5B**) without affecting its normal behavior and long-term survival (**Figure 5C**).

**Figure 5:**
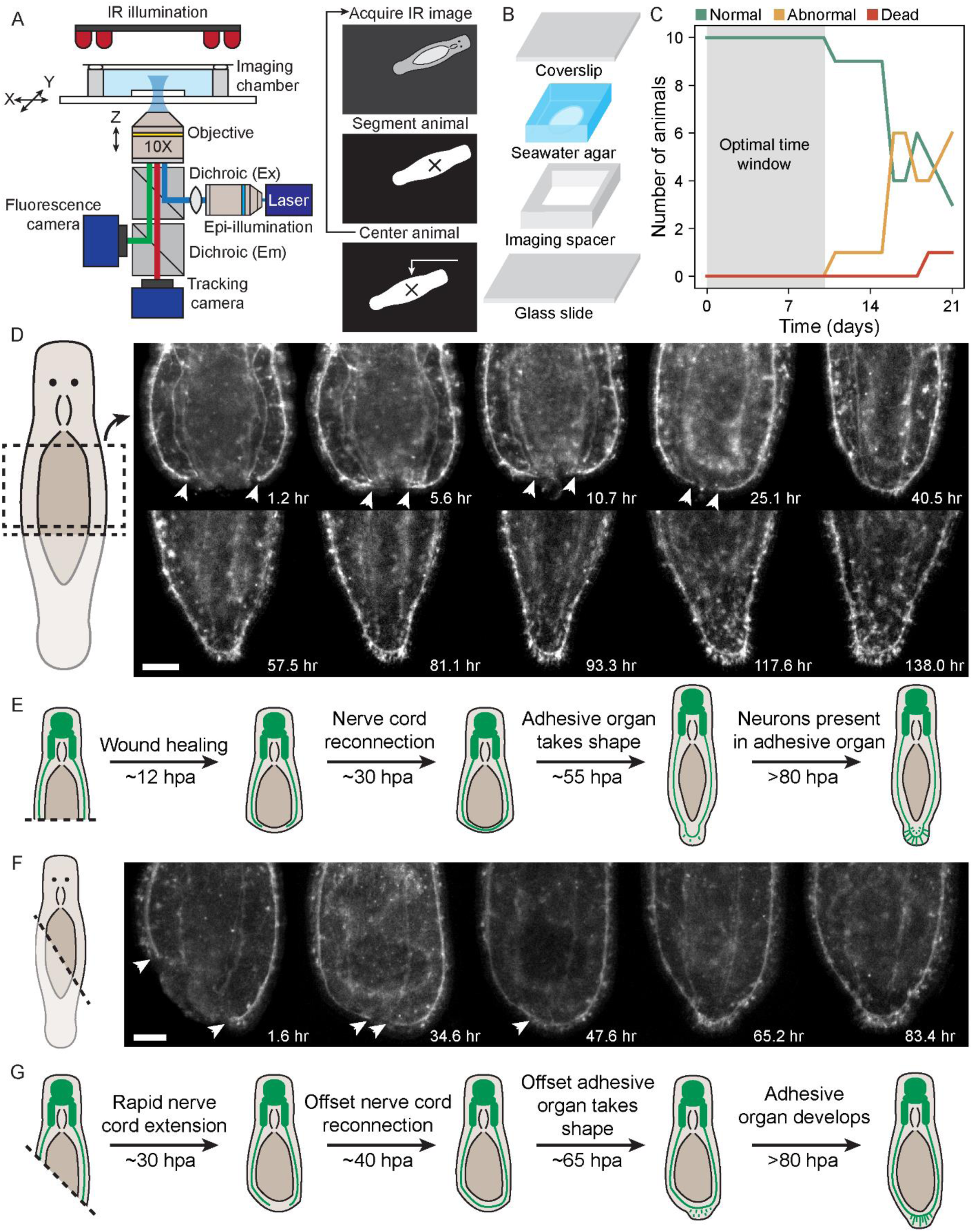
Tracking microscope enables continuous imaging of posterior neural regeneration in free-moving animals. **(A)** A diagram of the tracking microscope (left). An overview of the tracking routine involving infrared (IR) image capture, segmentation, and stage repositioning (right). **(B)** An exploded diagram of the long-term imaging chamber. **(C)** Survival curve of 10 animals placed in individual chambers and maintained at room temperature in the dark. All animals were phenotypically normal and actively moving up to 9 days, after which time half of the animals began limiting their movement and forming mucus cysts. By week three, only one animal had lysed. **(D)** Representative images of neural regeneration from a head fragment taken from a continuous week-long tracking microscopy session. Images are from the region highlighted in the cartoon (dashed box). Scale bar: 100 µm. **(E)** Cartoon showing the different stages of neural regeneration following a horizontal cut. By 12 hpa, the posterior tissue closes; 30 hpa, the ventral nerve cords reconnect; 55 hpa, additional neurons appear in the tail plate, 80 hpa, the tail continue to add new neurons while refining its shape. **(F)** Representative images of neural regeneration from an oblique cut. Arrows: termini of the ventral nerve cords. Scale bar: 100 µm. **(G)** Cartoon showing regeneration of the nervous system after an oblique cut. The oblique cut introduces an asymmetry evident in the uneven extension of the major nerve cords and offset adhesive organ. In the first 30 hpa, the left nerve cord extends a longer distance than the right, eventually meeting slightly off-center by 40 hpa. By 65 hpa, the tail plate begin to form adjacent the point of nerve cord reconnection, consistent with the anatomical posterior of the animal, and the tail continue to re-center by 80 dpa.

We horizontally bisected the PC2 strain and immediately mounted the head fragments for imaging. Initially, the gut protruded out of the wound and the nerve cords were severed, terminating abruptly at the injury site. Within the first 12 hrs, the gut retracted into the body, but the nerve cords remained disconnected. The two cords gradually extended towards each other and finally reconnected at ∼30 hpa. Once connected, the posterior tissue began to grow outwards, forming a new tail plate between 40 to 50 hpa. As the tail plate restored its characteristic pad shape, the nerve cords remained on the periphery of the newly regenerated tissue. Additional neurons started to emerge posterior to the nerve cords around 55 hpa, eventually arranging into their usual fan-like configuration in the adhesive organ by ∼100 hpa (**Figure 5D-E**, **Video S5**).

Intrigued by the bilateral symmetry observed during regeneration, we wondered how asymmetric amputation might affect this process. Specifically, would the nerve cords reconnect at the original midline or at the midpoint between the severed cords? To investigate this, we amputated animals at a 45-degree angle and monitored their regeneration using the tracking microscope. Initially, one of the severed nerve cords was ∼90 µm longer than the other. As the wound closed, the shorter nerve cord extended quickly towards the posterior while the longer nerve cord grew laterally a slight distance (**Figure 5F**). This asymmetry resulted in the two nerve cords meeting only slightly offset from the midline. By ∼65 hpa, neurons of the adhesive organ began forming at the midline adjacent to nerve cord closure site (**Figure 5G**, **Video S6**), suggesting that the specification of the adhesive organ is influenced by global body patterning cues, rather than the cues that instruct the location of nerve cord reconnection. To further confirm this impression, we amputated an animal at an extreme angle, which caused the remaining posterior tissue to wrap around the wound site, resulting in the nerve cords reconnecting on one side of the animal. Even in this extreme case, neurons associated with the tail plate regenerated further posterior than the site of nerve cord closure (**Figure S5A-B**), supporting that nerve cord closure and posterior regeneration may be controlled by distinct cues.

### Continuous live imaging allows quantification of regeneration progress across scales

With continuous tracking and live imaging, we now have the capability to simultaneously quantify the regeneration process across tissue, organismal, and behavioral scales in a single experiment. For example, at the tissue level, we annotated the nerve cords and measured their length following the 45-degree oblique cut. Strikingly, the shorter nerve cord extended linearly at a rate twice higher than the longer nerve cord (∼10 µm/hr vs. ∼5 µm/hr) (**Figure 6A**), indicating an adaptive mechanism where nerve cords modulate their extension rate in a manner reflecting the distance needed to travel for reconnection. Consistently, the distance between nerve cords also decreased linearly, after both oblique and symmetric cuts, though the absolute rates varied between animals and cuts (**Figure 6A**, **Figure S5C**).

**Figure 6:**
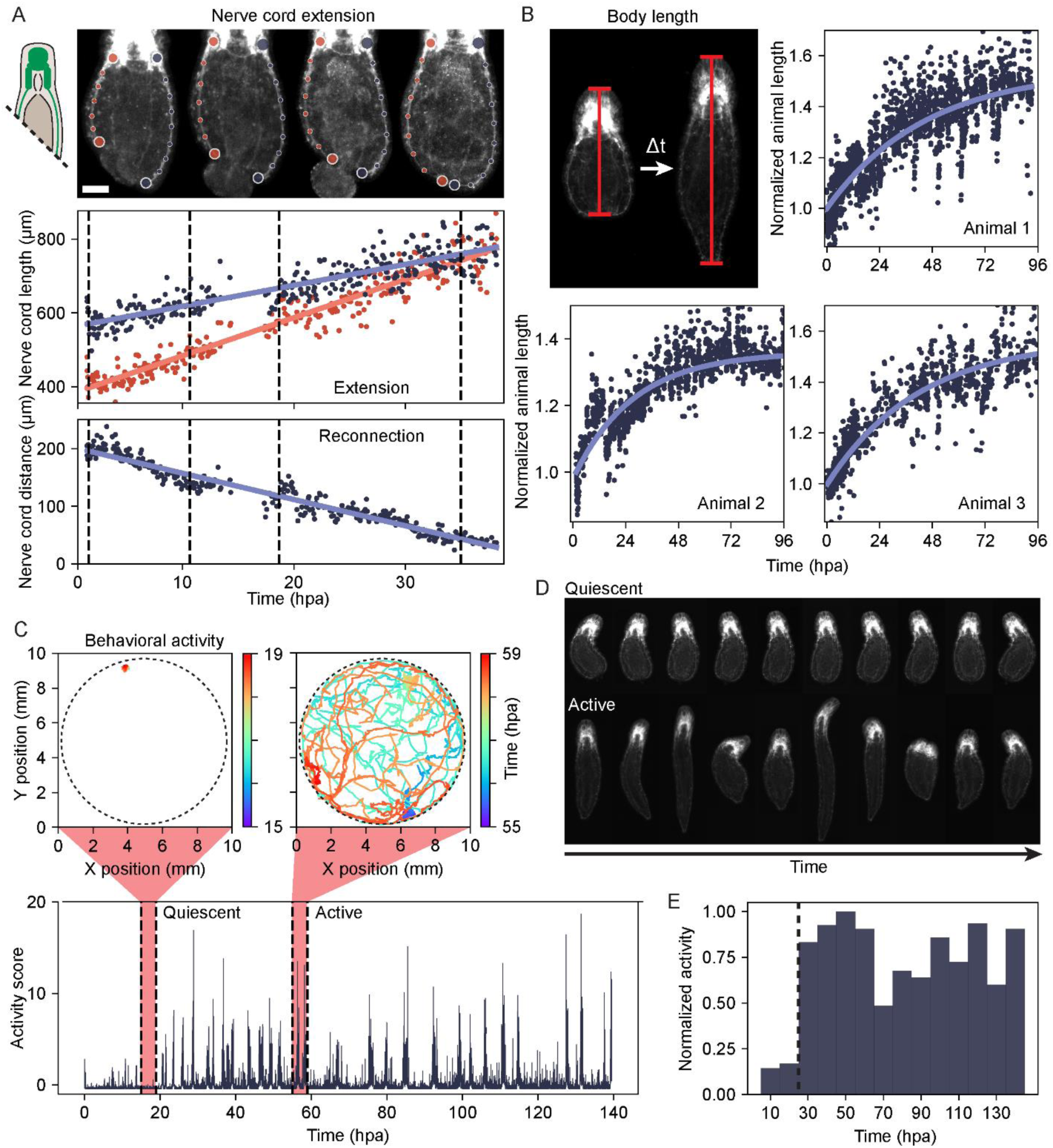
Continuous live imaging allows quantification of regeneration progress across scales. **(A)** A cartoon of oblique amputation (left). Nerve cord are manually annotated from the lateral ganglia (large upper dots) to the termini of the nerve cords (large lower dots). Scale bar: 50 µm. The length of each nerve cord is plotted over time showing linear extension (middle). The Euclidian distance between the nerve cord termini also decreases linearly (bottom). Dashed lines: the times corresponding to the representative images of the animal shown above. **(B)** An example showing body extension during regeneration (top left). Quantification of body length across 3 animals. The length of the animal is normalized to the starting length, and the data is fit to a curve *y* = *y*_0_ − *y*_1_*e*^−*kx*^ where *y* is the length of the animal, *y*_0_ is the final length, *y*_0_ − *y*_1_ is the starting length of the animal, *k* is the time constant, and *x* is time. Dots: data points. Lines: best fit. **(C)** The activity score of a tracked animal over ∼140 hrs. The highlighted regions correspond to a period of quiescence (left) in which the animal remains stationary, and a period of high activity (right) in which the animal explores much of its enclosure. Trajectories of the animal’s position over time in each time period are shown above. **(D)** Example images of tracked animals during the quiescent and active periods. Each image is spaced ∼20-25 min apart. During times of high activity, the animal is often stretched whereas the animal occasionally bends during the quiescent period. **(E)** The distribution of normalized average activity within 10 hr windows measured from the activity score in panel C. Dotted line: low average activity during the first 30 hpa, followed by an increase in average activity.

At the organismal level, the animals rescaled their bodies as regeneration progresses. Using an automated pipeline to segment and orient animals, we quantified their length throughout the course of regeneration. Unlike linear nerve cord extension, the animal length grew according to a ‘saturation curve,’ with the rate of regeneration proportional to the amount of tissue to be regenerated. Surprisingly, the time constant of this saturation appeared consistent across animals at 0.027±0.004 hr^-1^ (**Figure 6B**), suggesting a potentially characteristic time scale for *Macrostomum* tail regeneration.

At the behavioral level, the microscope’s tracking feature allowed us to quantify the animal movement with a time step of 2 s using the stage’s position and the animal’s location within the FOV. From the velocity data, we calculated a behavioral activity score (Bray et al., 2023), with high activity scores corresponding to animals actively exploring their environment, punctuated by body extension and scrunching (**Figure 6C-D**). After amputation, we observed a decrease in average activity during the first 20 hr, followed by a 50 hr bout of hyperactivity, eventually returning to the baseline level (**Figure 6E**). Correlating the behavioral output with the steps in tissue regeneration (**Figure 5D**), our data suggested that the animal’s activity was suppressed during wound healing. Overall, our tracking microscope represents a major advance in our capacity to simultaneously collect tissue and morphological data with high temporal resolution in freely moving animals throughout the regeneration process. This approach enables the quantitative analysis of neural regeneration while simultaneously providing a rich behavioral dataset to investigate how these processes are realted across scales.

### Multimodal imaging characterizes posterior regeneration defects induced by *β-catenin* RNAi

In planarian flatworms, knockdown of *β-catenin* causes anterior-posterior (A-P) patterning defects (Gurley et al., 2008; Petersen & Reddien, 2008), and in other systems, *β-catenin* is known to play important roles in neural induction and axonal regeneration (Rocheleau et al., 1999; Onishi et al., 2014; Watanabe et al., 2014; Garcia et al., 2018). This prompted us to postulate that *β-catenin* knockdown could block or incorrectly guide nerve cord extension post-injury in *M. lignano*.

To elucidate the role of *β-catenin* in posterior regeneration, we first sought a molecular posterior marker that reactivates early after tail amputation. In planarians, *wnt-1* marks the posterior pole, expressed in a single train of cells along the midline and is quickly induced at injury sites (Petersen & Reddien, 2009). We identified a *wnt-1* ortholog in the *M. lignano* genome (**Figure S6A**), which was upregulated in the posterior wound after amputation. We developed a hybridization chain reaction (HCR) protocol to detect *wnt-1* expression. In intact animals, *wnt-1* was expressed in a wide swathe of anchor cells located in the posterior adjacent to the adhesive glands stained by peanut agglutinin (PNA) (Lengerer et al., 2016) (**Figure 7A**). Following injury, *wnt-1* expression was induced within the posterior blastema (**Figure 7B**), providing a means to assess whether the posterior is successfully re-specified at the early stages of regeneration.

**Figure 7:**
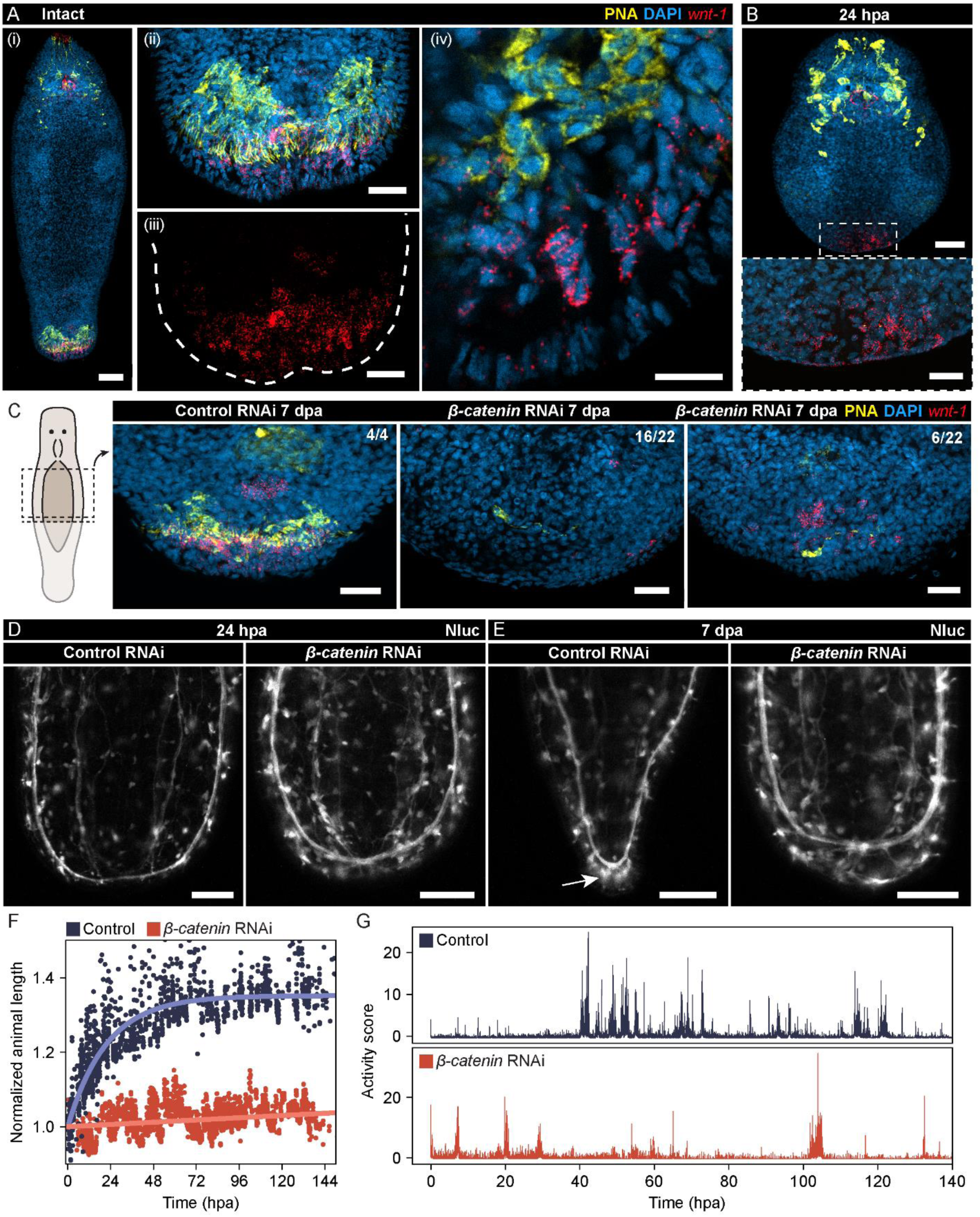
Multimodal imaging characterizes posterior regeneration defects in *β-catenin* RNAi animals. **(A)** Normal homeostatic appearance of the adhesive glands (yellow) and *wnt-1* expression (red) in the posterior. (i) An overview of the whole animal. (ii) A magnified view showing the *wnt-1*^+^ cells posterior to the adhesive glands. (iii) The same view as (ii) showing just *wnt-1* expression. Dashed line: outline of the tail plate. (iv) *wnt-1*^+^ anchor cells and PNA stained adhesive glands are adjacent to each other in the posterior. Scale bars: 50 µm (i); 20 µm (ii); 20 µm (iii); 10 µm (iv). **(B)** Normal animals showing *wnt-1* expression at 24 hpa. An overview of the animal (top), with a magnified view of the posterior (bottom) showing no adhesive glands but numerous *wnt-1*^+^ cells. Scale bar: 50 µm (top); 20 µm (bottom). **(C)** A cartoon of the amputation made (far left). The posteriors of control (left, n = 4/4) showing a regenerated array of adhesive glands and *wnt-1*^+^ anchor cells. In contrast, *β-catenin* RNAi animals show none (middle, n = 16/22) or few (right, n = 6/22) *wnt-1*^+^ cells and no adhesive glands at 7 dpa. Scale bars: 20 µm. **(D-E)** Luminescence imaging of control (left) and *β-catenin* RNAi (right) treated PC2 animals. At 24 hpa (D), animals in both groups healed the wound and the ventral nerve cords reconnected. At 7 dpa (E), control animals regenerated a full tail with neurons innervating the adhesive organ (arrow), *β-catenin* RNAi animals showed no tail regeneration or adhesive organ neurons radiating from the ventral nerve cord. Scale bars: 100 µm. **(F)** Normalized body length of control (blue) or *β-catenin* RNAi animals (red) showing now growth after *β-catenin* knockdown. Dots: individual data points. Lines: best fit. **(G)** Activity score of control animals (top) compared to *β-catenin* RNAi animals (bottom) showing reduced activity and the lack of recovery after *β-catenin* knockdown.

*M. lignano* has three *β-catenin* homologs, which we targeted with double stranded RNA (dsRNA) designed against a common sequence for RNAi-induced gene silencing (Mouton et al., 2023) (**Figure S6B**). Whereas control animals regenerated both adhesive glands and *wnt-1^+^* anchor cells, *β-catenin* RNAi animals failed to regenerate posterior structures, with *wnt-1^+^* cells absent in the majority (n=16/22) or rarely observed (n=6/22) (**Figure 7C**), indicating that regeneration was halted at an early stage under *β-catenin* knockdown conditions.

Remarkably, despite the impaired regeneration within *β-catenin* knockdowns, nerve cords still extended and reconnected normally by 24 hpa, as observed in the PC2 reporter strain through luminescence imaging (**Figure 7D**). Yet, beyond nerve cord reconnection, regeneration ceased (**Figure 7E**): tracking microscopy revealed no significant changes in the animal length over time, suggesting a complete lack of regenerative growth (**Figure 7F**). Furthermore, tracking data revealed a marked reduction in the behavioral activity of the *β-catenin* knockdown animals compared to controls in the process of regeneration (**Figure 7G**), indicating that nerve cord reconnection alone is insufficient to restore normal behavior. These findings are consistent with the notion that nerve cord reconnection is a part of the wound healing process, which can occur even after halting regeneration by *β-catenin* knockdown. Together, these results demonstrate the utility of our toolbox in dissecting complex regeneration phenotypes caused by genetic manipulations.

## Discussion

Whole-body regeneration is a complex, dynamic, organism-wide process involving many different cell types (Reddien, 2018; Fan et al., 2023) that has been challenging to study due to the limited tools available for transgenic manipulations and sensitive, longitudinal live imaging. Here, we presented three key techniques to accelerate the use of *M. lignano* as a platform system for studying whole-body regeneration. First, we established a modular genetic toolkit for rapidly assembling transgenes. As our research community matures, this toolkit will facilitate the engineering of increasingly sophisticated genetic constructs as labs contribute compatible parts to expand the library. Second, we demonstrated the use of chemical cellular ablation in a regenerative flatworm species, opening the door to investigating the regenerative potential of specific cell types and their individual contributions in the context of tissue regeneration. To complement targeted ablation, we developed an affordable and scalable open-source luminescence/fluorescence microscope to monitor the behavior of remaining cells post-ablation with high sensitivity, spatial resolution, and accuracy. Finally, we built a fluorescence tracking microscope, allowing for the observation of regeneration from initiation to completion, acquiring dynamic, quantitative data across multiple scales, ranging from the microscopic details within the tissue, through the macroscopic changes at the organismal level, and ultimately to the complex outputs of animal behavior. Collecting these multi-scale data on the same animal within a single experiment can help to understand how cellular and tissue-level processes translate into functional outcomes at both the organismal and behavioral levels.

By integrating these tools— transgenesis, ablation, and continuous live imaging – we pave the way for tagging and ablating various cell types (or even sub-types) to systematically delineate their roles during whole-body regeneration and to explore the crosstalk between these cell types. For example, our neural ablation experiments highlighted the critical role of the nervous system in facilitating wound healing. The tools are now becoming available to identify which neuronal populations are essential in this process, and whether neurons communicate directly with epidermal cells, as recently noted in the fly gut post-injury (Petsakou et al., 2023), or indirectly via other cell types, such as muscles or phagocyte-like cells we observed within the blastema.

A remarkable finding from our work is the differential extension rates of nerve cords following an oblique cut: the shorter nerve cord extended at a rate that was double that of the longer one. This suggests that the regulation of nerve cord repair may be adaptive, with the extension rate reflective of the distance needed to travel for reconnection. Interestingly, once the nerve cords began to extend, they extended at a constant rate until the nerve cords reconnected, indicating that the rate became fixed early in the repair process. This behavior contrasted with the growth of body length, which followed a saturation curve, meaning that the growth rate decreases as regeneration advances. Unraveling the mechanisms underlying this distinct kinetic pattern in neural repair is an important avenue for future research.

Intriguingly, the site at which nerve cords reconnected after oblique cuts did not necessarily correspond to the new posterior end of the body, where the adhesive organ consistently formed and where *wnt-1* expression was reliably reactivated. This observation, alongside the fact that nerve cords reconnected even after *β-catenin* RNAi – which resulted in the absence of posterior regeneration and the loss of posterior identity as indicated by *wnt-1* expression – suggests that nerve cord repair may operate independently of A-P patterning cues. This may explain how nerve cords can begin to extend before the new posterior gets re-specified. This is further supported by the ability of nerve cords to reconnect in the anterior of the non-regenerative tail fragments (**Figure S7A**). Possible mechanisms for guiding nerve cord extension and reconnection may be midline genes, known to be crucial for axonal guidance in various systems (Blockus & Chédotal, 2016), which may also function in conjunction with other morphogenic cues (Charron et al., 2003), as well as genes that may regulate mediolateral patterning, including the non-canonical Wnt signaling or planar cell polarity pathway (Almuedo-Castillo et al., 2011; Onishi et al., 2014). With tools including RNAi, tracking microscopy, and the neuronal reporter strain, *M. lignano* can become a new system to dissect the molecular logic underlying axonal guidance in the context of regeneration.

Finally, our characterization of the *β-catenin* RNAi phenotype in *M. lignano* unveiled significant variations in how the Wnt signaling pathway specifies the A-P axis during regeneration across different flatworm species. Unlike the planarian *Schmidtea mediterranea*, which regenerates a head in place of a tail after *β-catenin* RNAi (Gurley et al., 2008; Petersen & Reddien, 2008), *M. lignano* simply failed to regenerate anything. Furthermore, in planarian species that exhibit compromised anterior regeneration, *β-catenin* RNAi can often rescue this deficiency (Sikes & Newmark, 2013; Vila-Farré et al., 2023); however, we observed no such rescue in non-regenerative *M. lignano* tails (**Figure S7B-C**). Another key difference is observed in the expression of *wnt-1*. In *M. lignano*, *wnt-1* is expressed in a wide swathe of cells around the posterior end, which is in stark contrast to its expression in the planarian, where it is restricted to a single line of cells strictly at the posterior tip of the animal body. This difference is compounded by the finding that, while the injury-induced *wnt-1* expression is *β-catenin* independent in the planarian (Petersen & Reddien, 2009), *β-catenin* RNAi largely abolished *wnt-1* expression in *M. lignano* after amputation (**Figure 7C**) but not during homeostasis (**Figure S6C**), suggesting that distinct regulatory relationships between *wnt-1* and *β-catenin* may respond to injury in these two flatworms. Lastly, the relationship between axon guidance and body patterning may differ between the planarian and *M. lignano*. In *S. mediterranea*, it has been recently shown that the regeneration of guidepost cells is coupled with body axis re-specification following amputation, and these guidepost cells are essential for guiding the reconnection of visual axons across midline and to the central nervous system, a process downstream of body patterning (Scimone et al., 2020). In contrast, nerve cord reconnection in *M. lignano* seems to progress independent of A-P axis regeneration. Overall, these observed differences highlight the utility of studying *M. lignano* as a critical model for examining the requirements and evolutionary modifications in the role of Wnt signaling in controlling neural repair and primary body axis regeneration.

### Limitations of the study

While our current protocol has made injection easier, the fundamental limitation remains the low rates of germline transmission due to random integration (∼1%). In an effort to enhance transformation efficiency, we experimented with injecting recombinant Tol2 transposase (rTol2) mixed with transgenes flanked by Tol2 inverted repeats. To date, this approach has not yielded higher efficiencies compared to random integration. However, with future work in both transposon-based and targeted integration, our modular cloning toolkit should facilitate both methodological comparisons and screening of alternative integration methods, with the overall design ensuring compatibility between existing and new plasmids.

Regarding the microscopy techniques, using the same objective for both tracking and imaging in the tracking microscope introduces a trade-off between spatial resolution and the tracking consistency, as higher magnification reduces the FOV, thereby increasing the likelihood of losing track of the animal. Moreover, the current setup relies on wide-field epifluorescence for imaging which has the limitations of reduced signal to noise ratio due to out-of-focus light and disruption in tracking cellular structures when they move out of focus due to animal rotation or deformation. These issues may be addressed in future iterations by using separate objectives for tracking and imaging and incorporating optical sectioning and/or multifocal imaging technologies.

## Acknowledgements

We thank E Berezikov and J Wudarski for sharing the *M. lignano* NL-12 strain, plasmids, and the experimental protocols for animal husbandry, E Davies along with members in the Berezikov and Davies labs for stimulating discussion. This work is supported by a Stanford Bio-X Interdisciplinary Initiative seed grant (IIP11-40), an NSF EDGE grant (IOS-1923534), and a NIH grants 1R35GM138061 to BW and R35GM130366 to AZF.

## Author Contributions

Conceptualization: RNH, JG, AZF, BW; Methodology: RNH, HL, CC, RRB, ES, MP; Instrumentation: HL, CC, ES; Investigation: RNH, SV, RRB, JG; Writing – Original Draft: RNH, BW; Writing – Review & Editing: RNH, HL, CC, JG, AZF, BW; Supervision: AZF, BW; Funding acquisition: AZF, BW.

## Key Resources Table

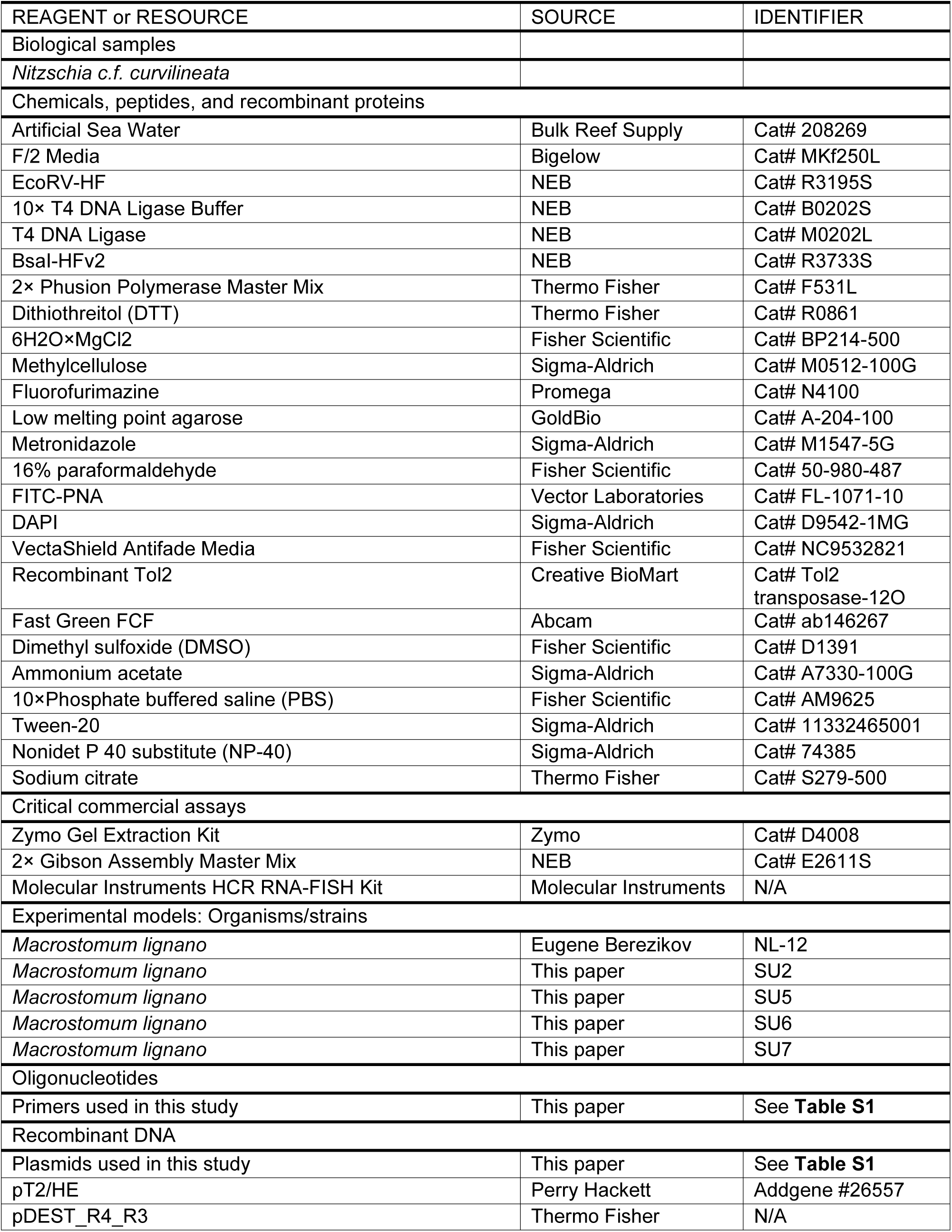
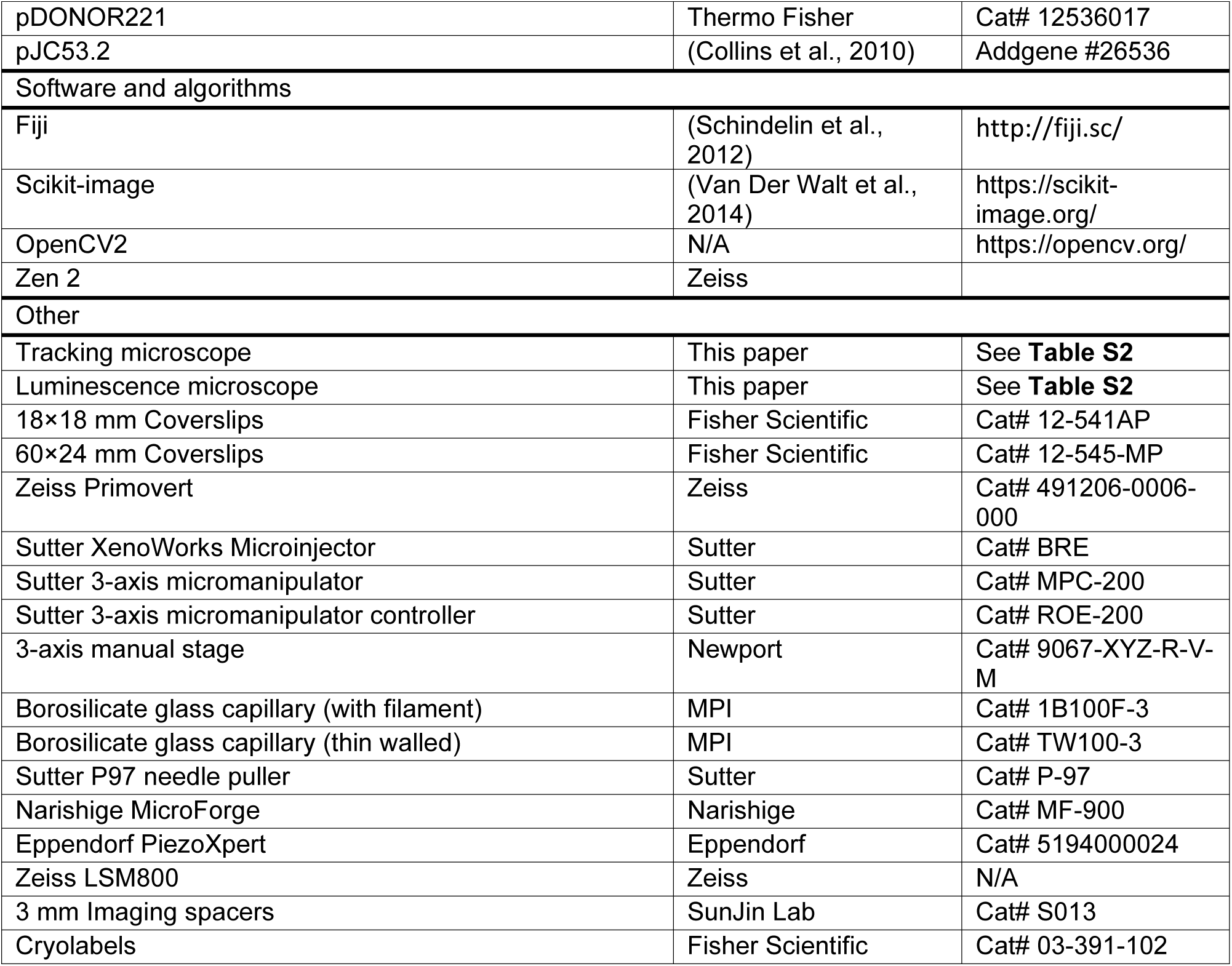

| REAGENT or RESOURCE | SOURCE | IDENTIFIER |
| --- | --- | --- |
| Biological samples |  |  |
| *Nitzschia c.f. curvilineata* |  |  |
| Chemicals, peptides, and recombinant proteins |  |  |
| Artificial Sea Water | Bulk Reef Supply | Cat# 208269 |
| F/2 Media | Bigelow | Cat# MKf250L |
| EcoRV-HF | NEB | Cat# R3195S |
| 10× T4 DNA Ligase Buffer | NEB | Cat# B0202S |
| T4 DNA Ligase | NEB | Cat# M0202L |
| Bsal-HFv2 | NEB | Cat# R3733S |
| 2× Phusion Polymerase Master Mix | Thermo Fisher | Cat# F531L |
| Dithiothreitol (DTT) | Thermo Fisher | Cat# R0861 |
| 6H<sub>2</sub>O×MgCl<sub>2</sub> | Fisher Scientific | Cat# BP214-500 |
| Methylcellulose | Sigma-Aldrich | Cat# M0512-100G |
| Fluorofurimazine | Promega | Cat# N4100 |
| Low melting point agarose | GoldBio | Cat# A-204-100 |
| Metronidazole | Sigma-Aldrich | Cat# M1547-5G |
| 16% paraformaldehyde | Fisher Scientific | Cat# 50-980-487 |
| FITC-PNA | Vector Laboratories | Cat# FL-1071-10 |
| DAPI | Sigma-Aldrich | Cat# D9542-1MG |
| VectaShield Antifade Media | Fisher Scientific | Cat# NC9532821 |
| Recombinant Tol2 | Creative BioMart | Cat# Tol2<br>transposase-120 |
| Fast Green FCF | Abcam | Cat# ab146267 |
| Dimethyl sulfoxide (DMSO) | Fisher Scientific | Cat# D1391 |
| Ammonium acetate | Sigma-Aldrich | Cat# A7330-100G |
| 10×Phosphate buffered saline (PBS) | Fisher Scientific | Cat# AM9625 |
| Tween-20 | Sigma-Aldrich | Cat# 11332465001 |
| Nonidet P 40 substitute (NP-40) | Sigma-Aldrich | Cat# 74385 |
| Sodium citrate | Thermo Fisher | Cat# S279-500 |
| Critical commercial assays |  |  |
| Zymo Gel Extraction Kit | Zymo | Cat# D4008 |
| 2× Gibson Assembly Master Mix | NEB | Cat# E2611S |
| Molecular Instruments HCR RNA-FISH Kit | Molecular Instruments | N/A |
| Experimental models: Organisms/strains |  |  |
| *Macrostomum lignano* | Eugene Berezikov | NL-12 |
| *Macrostomum lignano* | This paper | SU2 |
| *Macrostomum lignano* | This paper | SU5 |
| *Macrostomum lignano* | This paper | SU6 |
| *Macrostomum lignano* | This paper | SU7 |
| Oligonucleotides |  |  |
| Primers used in this study | This paper | See **Table S1** |
| Recombinant DNA |  |  |
| Plasmids used in this study | This paper | See **Table S1** |
| pT2/HE | Perry Hackett | Addgene #26557 |
| pDEST_R4_R3 | Thermo Fisher | N/A |
| pDONOR221 | Thermo Fisher | Cat# 12536017 |
| pJC53.2 | (Collins et al., 2010) | Addgene #26536 |
| Software and algorithms |  |  |
| Fiji | (Schindelin et al., 2012) | http://fiji.sc/ |
| Scikit-image | (Van Der Walt et al., 2014) | https://scikit-image.org/ |
| OpenCV2 | N/A | https://opencv.org/ |
| Zen 2 | Zeiss |  |
| Other |  |  |
| Tracking microscope | This paper | See **Table S2** |
| Luminescence microscope | This paper | See **Table S2** |
| 18×18 mm Coverslips | Fisher Scientific | Cat# 12-541AP |
| 60×24 mm Coverslips | Fisher Scientific | Cat# 12-545-MP |
| Zeiss Primovert | Zeiss | Cat# 491206-0006-000 |
| Sutter XenoWorks Microinjector | Sutter | Cat# BRE |
| Sutter 3-axis micromanipulator | Sutter | Cat# MPC-200 |
| Sutter 3-axis micromanipulator controller | Sutter | Cat# ROE-200 |
| 3-axis manual stage | Newport | Cat# 9067-XYZ-R-V-M |
| Borosilicate glass capillary (with filament) | MPI | Cat# 1B100F-3 |
| Borosilicate glass capillary (thin walled) | MPI | Cat# TW100-3 |
| Sutter P97 needle puller | Sutter | Cat# P-97 |
| Narishige MicroForge | Narishige | Cat# MF-900 |
| Eppendorf PiezoXpert | Eppendorf | Cat# 5194000024 |
| Zeiss LSM800 | Zeiss | N/A |
| 3 mm Imaging spacers | SunJin Lab | Cat# S013 |
| Cryolabels | Fisher Scientific | Cat# 03-391-102 |

## Resource availability

### Lead contact

Further information and requests for resources and reagents should be directed toward and will be fulfilled by the lead contact, Bo Wang.

### Materials availability

Plasmids used in this study are available upon request and have been deposited to Addgene (#IDs). Fluorescent/luminescent strains generated are available upon request.

### Data and code availability

Imaging and tracking data reported in this paper will be shared by the lead contact upon request. This paper does not report any original code.

## Experimental model and subject details

### Animal care and maintenance

*M. lignano* strain NL-12 (Wudarski et al., 2020) and derived strains were maintained at 20℃ and 60% humidity with a 14/10 day/night cycle in ASW (Bulk Reef Supply, Cat# 208269) with a specific gravity of 0.026. The diatom *Nitzschia c.f. curvilineata* was seeded in 150 mm glass dishes and grown in F/2 media (Bigelow, Cat# MKf250L). When confluent and ready for feeding to animals, the F/2 media was poured off and replaced with fresh ASW into which animals were transferred. Feeding was performed once a week. To maintain cultures suitable for egg preparation, age synchronized cultures were collected by removing adults from a plate after feeding and allowing the eggs to hatch. The juveniles from multiple plates were pooled and used to seed a fresh plate of age synchronized animals.

## Method Details

### Cloning

All primers and oligos used are referred to in **Table S1**, and all parts and constructs have been included in **File S1**. Parts (promoters, genes, terminators) were assembled by PCR amplification of the desired sequence with overhangs including Gibson assembly homology arms and BsaI restriction sites. The sequences are listed in **Table S1**

GeNL (Suzuki et al., 2016) and NTR2.0 (Sharrock et al., 2022) were codon optimized using https://www.macgenome.org/codons/ without introns and synthesized by Twist Bioscience. GeNL-P2A-NTR2.0 was assembled by Gibson assembly. Promoters from *enolase* and *pc2* were amplified from genomic DNA. mScarlet as well as promoters from *eef1α, apob,* and *myh6* were amplified from plasmids described in (Wudarski et al., 2017).

pEmpty was linearized with EcoRV-HF (NEB, Cat# R3195S), run on a 1% agarose gel, and extracted using the Zymo Gel Extraction kit (Zymo, Cat# D4008). Gibson assembly was performed by combining a part amplicon and linearized pEmpty in a 2:1 mass ratio in 2× NEB Gibson Assembly Master Mix (NEB, Cat# E2611S) and incubating at 50℃ for 2 hr before transformation. Clones were sequenced using M13 Forward (5’-GTAAAACGACGGCCAGT-3’) or M13 Reverse (5’-CAGGAAACAGCTATGAC-3’) primers.

Unique sequence (UNS) oligos were annealed by combining sense and antisense oligos at 5 µM in duplex buffer (IDT, Cat# 11-05-01-12), incubated at 94 ℃ for 2 min, and cooled slowly at a rate of 1 ℃/min. The duplexed oligos where then diluted to 50 nM in H_2_O.

Parts were assembled into TUs via Golden Gate assembly. 0.5 µL of each part (promoter, gene, and terminator, 30 nM), 0.5 µL of 5’ and 3’ UNS oligos each (50 nM) corresponding to their position in the final plasmid, 0.5 µL of 10× T4 DNA Ligase Buffer (NEB, Cat# B0202S), 0.25 µL of T4 DNA Ligase (NEB, Cat# M0202L), and 0.25 µL of BsaI-HFv2 (NEB, Cat# R3733S), were mixed to a final volume of 10 µL in nuclease free water. The samples were cycled with the following program: [37 ℃ for 3 min, 16℃ for 4 min] × 12, 50℃ for 5 min, hold at 4℃.

For example, a single TU vector would require only UNS1A and UNS10D oligos in the prior Golden Gate reaction and would be amplified using UNS1F and UNS10R primers. Alternatively, dual TU vectors would require the first TU use UNS1A and UNS3D oligos, while the second would require UNS3A and UNS10D oligos, and would be amplified by UNS1F/UNS3R, and UNS3F/UNS10R primers, respectively.

Assembled TUs were amplified by PCR from the prior Golden Gate reaction products. 2 µL of Golden Gate reaction were added to 25 µL of 2× Phusion Polymerase Master Mix (ThermoFisher, Cat# F531L), mixed with 18 µL of H_2_O, and 3 µL of 10 µM forward and reverse primers corresponding to the terminal UNS sequences used for that TU. The samples were cycled with the following program: 98℃ for 30 sec, [98℃ for 30 sec, 60℃ for 30 sec, 72℃ for 30 sec per kb] × 35-40, 72℃ for 5 min, hold at 4℃. The resulting product was run on a 1% agarose gel and the band corresponding to the TU was extracted using the Zymo Gel Extraction kit and eluted in 11 µL of H_2_O.

pDest and pTol2Dest were constructed by performing primer extension PCR and Gibson assembly to insert an EcoRV-HF restriction site between a 5’ UNS1A and 3’ UNS10D sequences. pDest was constructed by amplification of pDEST_R4-R3 (Invitrogen) with primers BW-NH-710 and BW-NH-711 while pTol2Dest was constructed by amplification of pT2/HE (Addgene Plasmid ID: 26557) with BW-NH-792 and BW-NH-793. The backbone amplicons were then self-ligated using Gibson assembly. Unlike pDest, pTol2Dest contains terminal repeats for integration via Tol2 transposase.

TUs were assembled into the destination vectors, pDest or pTol2Dest, via Gibson assembly. Destination vectors were linearized using EcoRV-HF, run on a 1% agarose gel, and extracted using the Zymo Gel Extraction kit. Gibson assembly was performed combining TUs and linearized destination vector in a 2:1 mass ratio in 2× NEB Gibson Assembly Master Mix. The samples were incubated at 50℃ for 2 hr before transformation.

### Egg preparation

To prepare fertilized eggs for microinjection, ∼200-300 individual gravid animals (evident by a large, well-developed oocyte present in their posterior) from age-synchronized populations at ∼4-7 days post feeding were collected into two separate 60 mm dishes filled with ASW. Every hour, freshly laid eggs were transferred using an eyelash pick and transferred to the up-side down lid of a 100 mm polystyrene petri dish, which was cooled to 4 ℃ to halt the development at 1-cell stage. Once ∼100-150 eggs were lined up on the lid, the eggs were washed in 30 mM DTT in ASW for 5-10 min, swirling the plate occasionally. The eggs were then washed three times with ASW before microinjection.

### Microinjection

Eggs were injected on a Zeiss Primovert (Zeiss, Cat# 491206-0006-000) inverted stereomicroscope equipped with a Sutter XenoWorks electronic micromanipulator (Sutter, Cat# MPC-200), controller (Sutter, Cat# ROE-200) and pressure source (Sutter, Cat# BRE). A manual 3-axis stage was used to manipulate the holding pipette (Newport, Cat# 9067-XYZ-R-V-M). Needles were pulled from borosilicate glass capillaries (MPI, Cat# 1B100F-3) on a Sutter P97 needle puller using a 3-step pulling protocol (1. Heat = 754, Pull = 90, Velocity = 8, Time = 250; 2. Same as 1. 3. Heat = 754, Pull = 85, Velocity = 8, Time = 250). A 20-degree bend at the tip of the needle was introduced using a Narishige MicroForge (Narishige, Cat# MF-900). Holding pipettes were pulled from borosilicate glass capillaries (MPI, Cat# TW100-3) (Heat = 738, Pull = None, Velocity = 150, Time = None) and cut, flame polished, and bent to a 30-degree angle.

Injection mix was prepared by adding 1 µL of recombinant Tol2 transposase (rTol2, Creative BioMart) and 0.5 µL of Fast Green FCF (Abcam, Cat# ab146267) followed by plasmid DNA and rTol2 storage buffer (10 mM HEPES, 300 mM KCl, pH 6.9) to a final plasmid concentration of 50 ng/µL. Injection in the absence of rTol2, however, yielded similar transformation efficiencies. The solution was mixed and centrifuged for 1 min to pellet any small particles that may clog the needle. Injection mix was then loaded by back filling the needle. Injections were performed using an initial pressure of 1,000 hPa and back-pressure of +100 hPa, which was adjusted depending upon the flow of the needle. Without applying negative pressure, the holding pipette was placed behind the egg as a backstop. A PiezoXpert (Eppendorf, Cat# 5194000024) was used to assist in penetrating the oocyte’s membrane with settings “Int = 86, Speed = 10, Pulse”. Mix was ejected until a visible bolus of fluid became visible within the oocyte.

### Amputations

Animals were anesthetized in 7.14% 6H_2_O×MgCl_2_ (Fisher, Cat# BP214-500) for 5 min and then amputated using a stainless-steel scalpel. Fragments were transferred to ASW to recover. Once mobile, fragments were transferred to chambers with fresh ASW for downstream analyses.

### Luminescence/fluorescence microscope

A thermoelectrically cooled camera using the Sony IMX571 sensor (ToupTek Cat# ITR3CMOS26000KMA) was chosen for its high quantum efficiency (91%), low read noise (<1.5e-at 12 dB gain), and low dark current. To increase photon capture efficiency, a 10 MP, f = 50 mm imaging lens was used as a tube lens to provide demagnification (enabling the use of higher NA objectives). The light tight enclosure was constructed from opaque black acrylic, laser cut, and assembled into a box with electrical tape to prevent light bleeding through the seams (**Figure S4A**). Existing and new Squid components were designed for optomechanical integration, with CAD files available at https://squid-imaging.org.

### Tracking microscope

Components comprising the microscope were constructed from modules described in (Li et al., 2020) and https://squid-imaging.org. Animals were illuminated with an IR LED light source at 850 nm. IR images for tracking were acquired at ∼6 Hz using a monochrome camera (Daheng Imaging, Cat# MER-1220-32U3M) at 10× magnification (BoliOptics, Cat# 03033331). Images were binarized by user-input threshold in the GUI. The binarized images were eroded and dilated to remove noise and fill gaps. OpenCV was used to calculate the centroid of the largest contiguous region. Centroid tracking was performed using a nearest neighbor approach in which the animal’s centroid in the subsequent frame was determined within a search radius centered on the animal’s previous centroid location. The displacement of the centroid was converted into x-y stage movements using a proportional-integral-derivative (PID) controller, implemented on an Arduino Due microcontroller. The stage position was then adjusted by a stepper motor with optical encoder to precisely maintain the centroid (and animal) in the center of the FOV. Code for tracking was adapted from https://github.com/prakashlab/squid-tracking.

While tracking, fluorescence images were acquired every 50 s. Fiber-coupled 405/488/561/638 nm lasers were despeckled through a Molex despeckler and used as excitation. A quad-bandpass dichroic filter set (405/488/561/640) was used to split the emission light (**Figure S4B**). Image acquisition software can be found at https://github.com/hongquanli/octopi-research.

### Live luminescence and fluorescence confocal imaging

Animals were first anesthetized in 7.14% 6H_2_O×MgCl_2_ and 2% methylcellulose (Sigma, Cat# M0512-100G) in deionized water and then transferred to a coverslip slide in a 20 µL droplet. For luminescence imaging, 0.5 µL of 8.7 mM Fluorofurimazine (FFz) (Promega, Cat# N4100) in PBS was added to the droplet and mixed thoroughly. The corners of a coverslip were then scraped across clay to make four small clay ‘feet’ and gently placed over the droplet. Using forceps, corners of the coverslip were pressed down to firmly restrict the animal in place. If the animal continued to move, a paper towel was used to wick away excess water from the underside of the slide to decrease the distance between the coverslip and the slide. The slide was then imaged on the luminescence microscope using an Olympus 20× objective (NA=0.75) with an exposure time varying between 10 and 60 s. For confocal imaging, slides were prepared as above, skipping the FFz step. Imaging was performed on a Zeiss LSM800 AxioObserver using a 40× water-immersion objective (NA=1.1). After short imaging sessions (∼10-20 min), animals may be recovered by adding a droplet of ASW to the corner of the coverslip and gently lifting it with a razor blade.

### Preparation of long-term imaging chambers

Animals were starved for 48 hr to reduce gut autofluorescence. Imaging chambers were produced by gluing a 3 mm imaging spacer (SunJin Lab, Cat# S013) to a glass slide and sticking 1-2 stacked circular cryolabels (Fisher Scientific, Cat# 03-391-102) in the center of the spacer. ∼2 mL of 2% low melting-point agarose (GoldBio, Cat# A-204-100) in ASW was pipetted into the imaging spacer until it completely fills the chamber. A second slide was pressed flat over the agarose carefully not to trap any bubbles. The agarose block was solidified in the fridge. A scalpel was used to separate the agarose block from the sides of the imaging spacer, and the block was placed upside down on a slide. A single animal was placed in 1-2% methylcellulose in ASW and then pipetted onto the depression in the agarose made by the cryolabels. A second slide with a 3 mm imaging spacer (without cryolabels) was then slowly lowered upside down onto the agarose block without generating any bubbles or removing the animal. Once placed, the whole assembly was flipped back right side up to remove the top glass slide. Finally, vacuum grease was applied to the top of the imaging spacer and a coverslip was placed over the top to seal the contents within the imaging spacer.

### Chemical ablation

MTZ (Sigma-Aldrich, Cat# M1547-5G) was dissolved in DMSO to a stock concentration of 1 M. Animals starved for 2 d and placed in 3 mL of ASW containing either 0.5 mM (gut ablation) or 5 mM MTZ (neural and muscle ablation). The ASW-MTZ mixture was pipetted up and down until there were no visible crystals of MTZ remaining. Controls contained an equivalent percentage of DMSO. ASW and MTZ was replaced every other day for the duration of ablation.

### RNAi

Primers (**Table S1**) were used to amplify a ∼500 bp region of *β-catenin* and TA-cloned into pJC53.2 (Addgene Plasmid ID: 26536) (Collins et al., 2010). Linear templates flanked by T7 promoters were generated using PCR. *In vitro* RNA synthesis was performed using T7 polymerase, RNA was precipitated using ammonium acetate (NH_4_Ac, 10 M) and ethanol, denatured, then re-annealed. Control RNAi was derived from the ccdB insert of the unmodified pJC53.2 plasmid.

RNAi was performed by soaking. Animals were starved overnight to eliminate diatoms from their gut. In a 24-well plate, ∼15-20 animals were placed in 1 mL of ASW containing 2 µg of dsRNA. The ASW and dsRNA mixture was replaced every other day for 3 weeks until amputation.

### Fixation

Animals were starved overnight in fresh ASW. Before fixation, animals were washed twice with ASW for 5 min each. ASW was then replaced with 2 mL of 7.14% 6H_2_O×MgCl_2_ for 5 min to relax the animals. 500 µL of the MgCl_2_ solution was removed and 500 µL of 16% paraformaldehyde (PFA) (Fisher Scientific, Cat# 50-980-487) was added to a final concentration of 4% PFA. After 15 min of fixation, 100 µL of 10% NP-40 was added. After 45 min of fixation, the fixative was replaced with PBS containing 0.1% Tween-20 (PBSTw). Animals were then washed twice with PBSTw for 5 min each, then dehydrated by incubating for 5 min in increasing concentrations of methanol (25%, 50%, 75%, 100%), which can be stored at -20℃.

### Hybridization chain reaction

Fixed animals in methanol were rehydrated by 5 min incubations in increasing concentrations of PBSTw (25%, 50%, 75%, 100%). Animals were incubated in a 1:1 volume ratio of PBSTw to probe hybridization buffer (Molecular Instruments) for 10 min at room temperature (RT). The solution was then replaced with pre-hybridization solution (Molecular Instruments) for 1 hr at 37 ℃. The pre-hybridization solution was then replaced with probe solution (4 pmol of probe mixture for every 500 µL of probe hybridization buffer). Probes targeting *wnt-1* were designed using https://github.com/rwnull/insitu_probe_generator and ordered from IDT. Probe sequences are included in **Table S1**.

Samples were incubated overnight (∼20 hr) at 37℃. Probe solution was removed, and the samples were washed 4 times with 500 µL of probe wash buffer (Molecular Instruments), pre-heated to 37 ℃, for 20 min each at 37 ℃, and then washed twice for 5 min each with 5× SSC supplemented with 0.1% Tween-20 (5× SSCT). Pre-amplification was performed by incubating samples with 500 µL amplification buffer (Molecular Instruments) for 30 min at RT. 30 pmol of hairpin H1 and 30 pmol of hairpin H2 in 10 µL of 3 µM stock solutions were snap cooled (heated to 95℃ for 90 sec and cooled to RT in a dark drawer for 30 min) separately for every 500 µL of amplification buffer. Snap cooled hairpins H1 and H2 (10 µL each) were added to a tube of 500 µL amplification buffer. Amplification buffer was removed from the samples and replaced with the hairpin-containing amplification buffer and incubated overnight (∼12-16 hr) in the dark at RT. On the following day, excess hairpins were removed by washing the samples with 500 µL of 5×SSCT at RT (2× for 5 min, 2× for 30 min, and finally once for 5 min). The samples were transferred to PBSTw by incubating in increasing concentrations of PBSTw in 5×SSCT (25%, 50%, 75%, 100%) for 5 min each. For visualizing the adhesive organs, samples were incubated in 1:200 FITC-PNA (Vector Laboratories, Cat# FL-1071-10) and 1:5000 DAPI (10 mg/mL, Sigma-Aldrich, Cat# D9542-1MG) for 30 min. Samples were washed 2× with PBSTw for 5 min each and mounted in VectaShield Antifade (Fisher Scientific, Cat# NC9532821) mounting solution and stored at 4 ℃.

## Quantification and Statistical Analysis

### Image processing

Confocal images were processed in the Zeiss Zen 2 software. Confocal stacks were aligned using the z-stack alignment tool. Dual-color images were spectrally de-mixed. Epifluorescence and luminescence images were processed using ImageJ v1.53k.

### Tracking microscope image processing

Raw images were processed using Python 3.8, Scikit-Image and OpenCV2. Images were first binarized by gaussian thresholding. An ellipse was fitted to the resulting segmented shape and the image was rotated to align the major axis of the ellipse vertically. The image was then rotated another 180° if the animal was detected with the anterior facing downward. Images were then cropped based on a common bounding box which encompassed the animal across all frames of the video. Nerve cords were manually annotated using a custom GUI for loading images and selecting anatomical landmarks. The body length of the animal was determined by the longest projection among the radially projecting rays every 5° from the centroid of the animal. Activity scores were calculated by denoising the instantaneous velocity, using a Morlet wavelet transformation and summing scales 1-31 (Bray et al., 2023). The activity scores are baseline subtracted.

## Supplemental Figures

**Figure S1:**
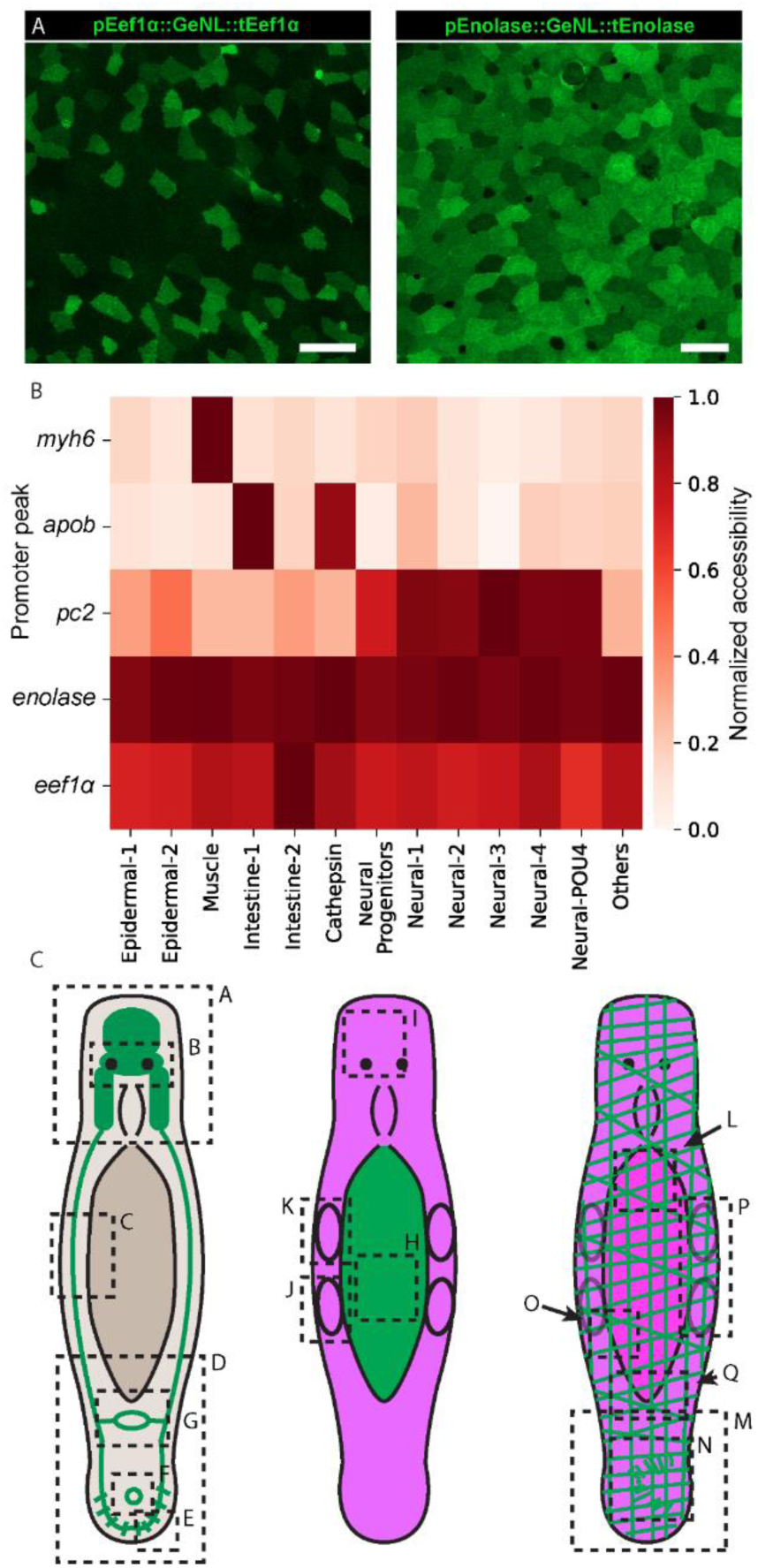
Reporter expression is consistent with single-cell gene expression analyses. Related to Figure 1 and 2. (**A**) Confocal images of live, anesthetized animals showing transgene expression in the epidermis. Expression is more stochastic when the reporter is driven by the Eef1α promoter (left) than the Enolase promoter (right). Green: mNeonGreen. Scale bars: 20 µm. (**B**) Single-cell ATAC-seq promoter peak accessibility for each promoter used in this study. The data is from (Chai et al., 2024). Briefly, *myh6* accessibility is specific to muscles; *apob* is accessible in a subpopulation of intestinal cells (intestine 1) and a subset of the *cathepsin^+^* phagocytic cells; *pc2* promoter is accessible in neural progenitors and all neural clusters; *enolase* shows uniform accessibility across all tissues, and *eef1α* has broad accessibility, with uniquely high expression in the intestine 2 cluster. (**C**) Diagrams of the three transgenic strains (PC2, APOB, MYH6. Dotted boxes correspond the region imaged in Figure 2 and its matching panel lettering (Figure 2A-Q).

**Figure S2:**
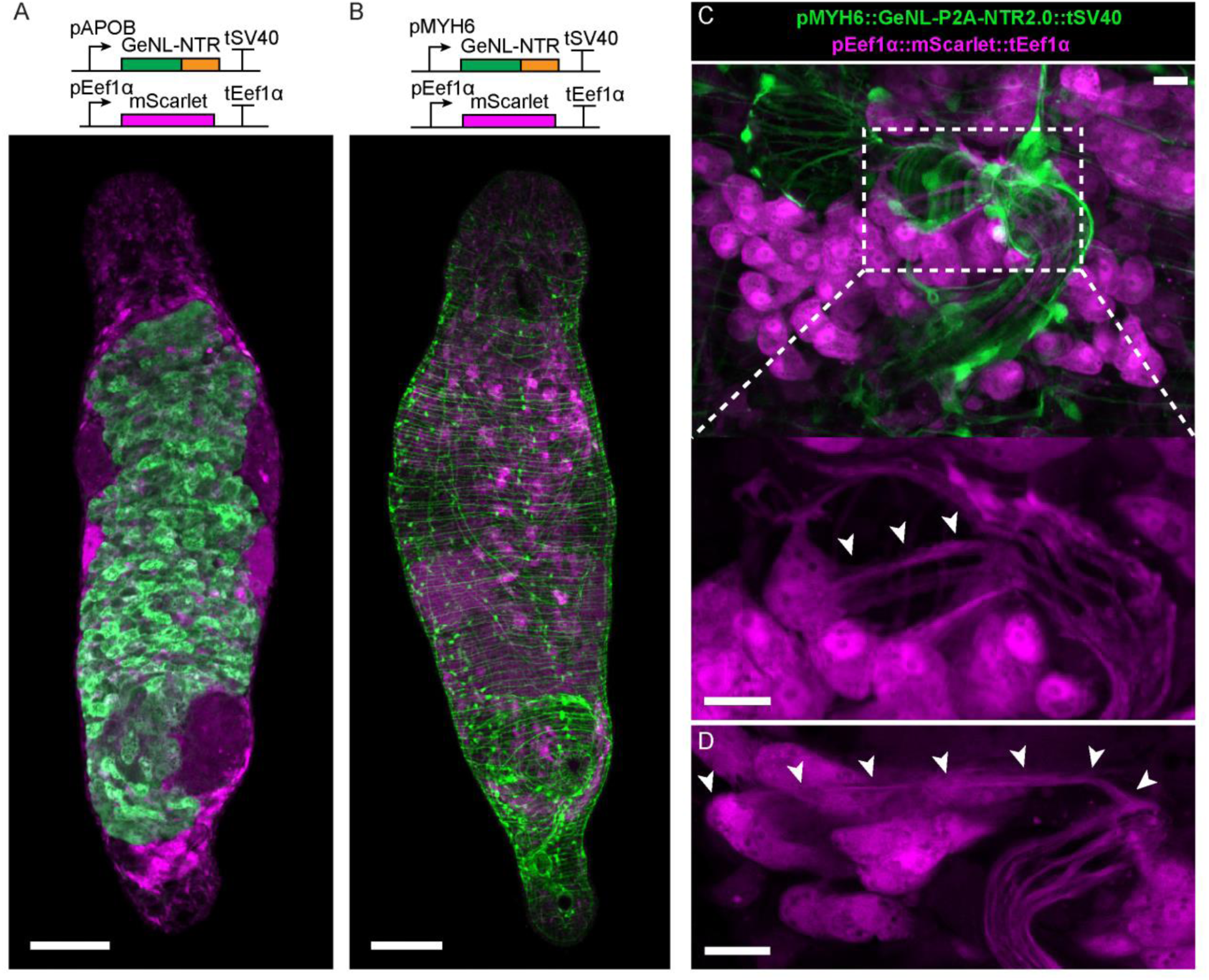
Dual-color reporters enable detailed morphological analysis. Related to Figure 2. (**A-B**) Confocal images showing mNeonGreen (green) and mScarlet (magenta) fluorescence in the pAPOB::GeNL-P2A-NTR2.0::tSV40, pEef1α::mScarlet::tEef1α (A) and in the pMYH6::GeNL-P2A-NTR2.0::tSV40, pEef1α::mScarlet::tEef1α reporter strains. Scale bars: 100 µm. (**C**) Confocal images showing the male copulatory apparatus. Inset: mScarlet expression showing prostate gland cells (magenta) extending processes into the muscle-bound copulatory system (dashed box). Arrows: the process. Scale bars: 10 µm. (**D**) Image of a prostate gland cell extending a long process multiple cell bodies across to invade the stylet. Arrows: the process. Scale bar: 10 µm.

**Figure S3:**
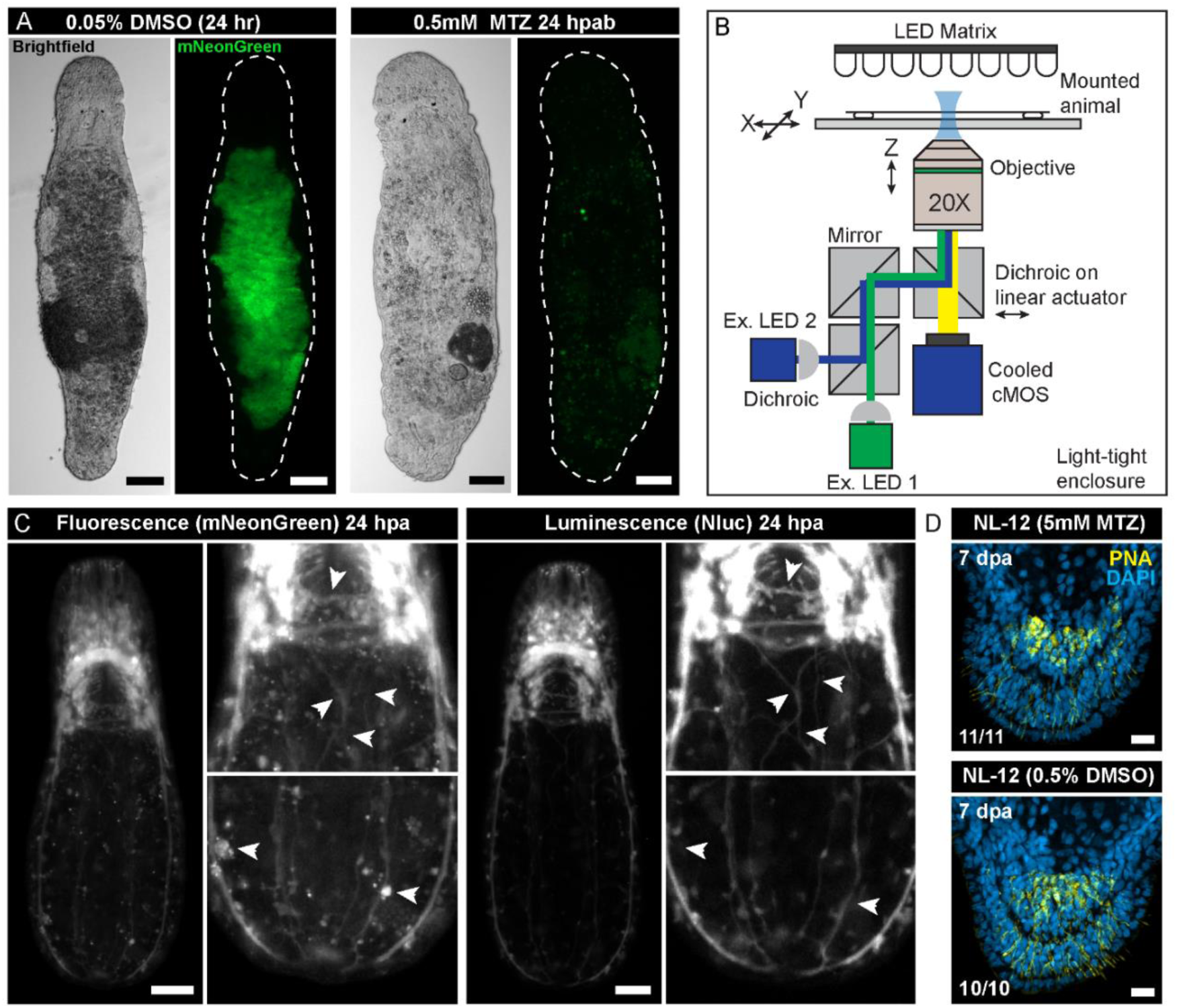
Fluorescence and luminescence characterization of the ablation strains. Related to Figure 4. **(A)** Gut ablation during homeostasis. Animals treated with DMSO (0.05%) show bright reporter signal in their gut after 24 hr of incubation (left). In contrast, animals incubated in MTZ (0.5 mM) showed no detectable gut fluorescence after 24 hr of treatment, suggesting efficient ablation (right). Scale bars: 100 µm. **(B)** Schematic showing the luminescence microscope design. Briefly, the animal is illuminated by an LED array for brightfield imaging. Two LEDs provide excitation at 375 nm, and 470 nm, respectively. A dichroic filter cube sits on a linear actuator controlling the filter switch. A cooled CMOS sensor detects both luminescence and fluorescence signal. The entire microscope is enclosed in a light-tight, temperature-controlled chamber. The motorized stage is controlled from a joystick and computer outside of the enclosure. All parts are listed in **Table S2**. **(C)** A comparison of fluorescence (left) and luminescence (right) images on the same animal 24 hpa. The magnified views show the anterior and posterior regions. Arrows: features for comparisons between fluorescence and luminescence images. Luminescence provides higher contrast of neural processes (top right) and eliminates autofluorescent puncta (bottom right). Scale bar: 50 µm. **(D)** Confocal images of wild-type (NL-12) animals treated for 6 d with MTZ (5 mM) (n = 11/11) or DMSO (0.5%) (n = 10/10) show no defects in posterior regeneration at 7 dpa while recovering in ASW, evidenced by the presence of PNA stained adhesive glands (yellow). Scale bar: 10 µm.

**Figure S4:**
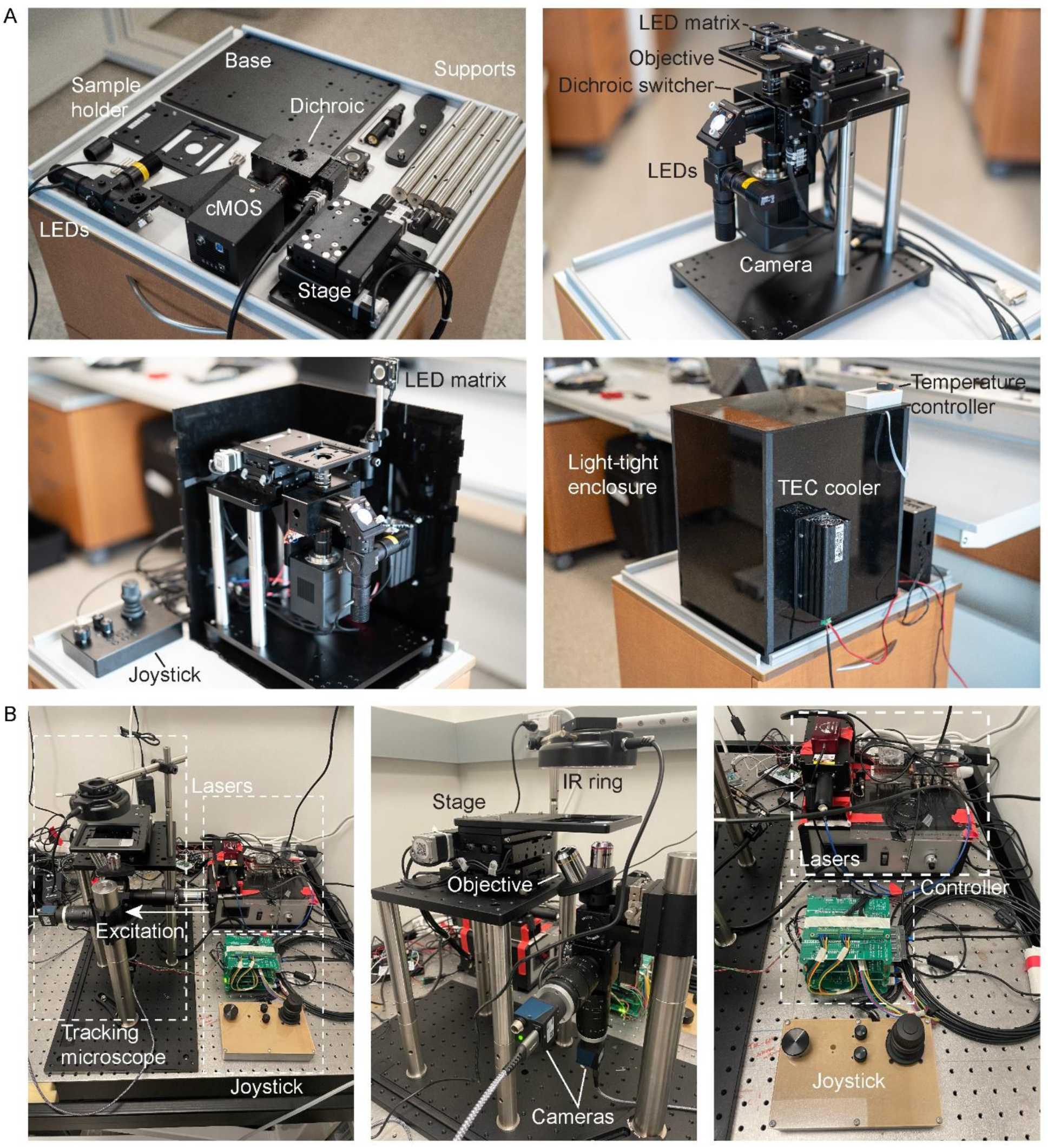
The luminescence/fluorescence and tracking microscopes. Related to Methods. **(A)** The components required to construct the luminescence microscope laid out (top left). The assembled microscope outside of its light-tight enclosure (top right). The microscope is situated within the light-tight enclosure (bottom left). The microscope is fully enclosed, showing the temperature controller incorporated into the enclosure (bottom right). Relevant components are labeled. **(B)** The tracking microscope consists of the body of the microscope, fiber coupled lasers and driver providing excitation light from the right, and joystick and stage controller (left). A ring of IR LEDs (850 nm) illuminates the sample from above. IR and fluorescence light is sent to separate cameras (middle). The microscope can be controlled manually with a joystick and through software while tracking (right).

**Figure S5:**
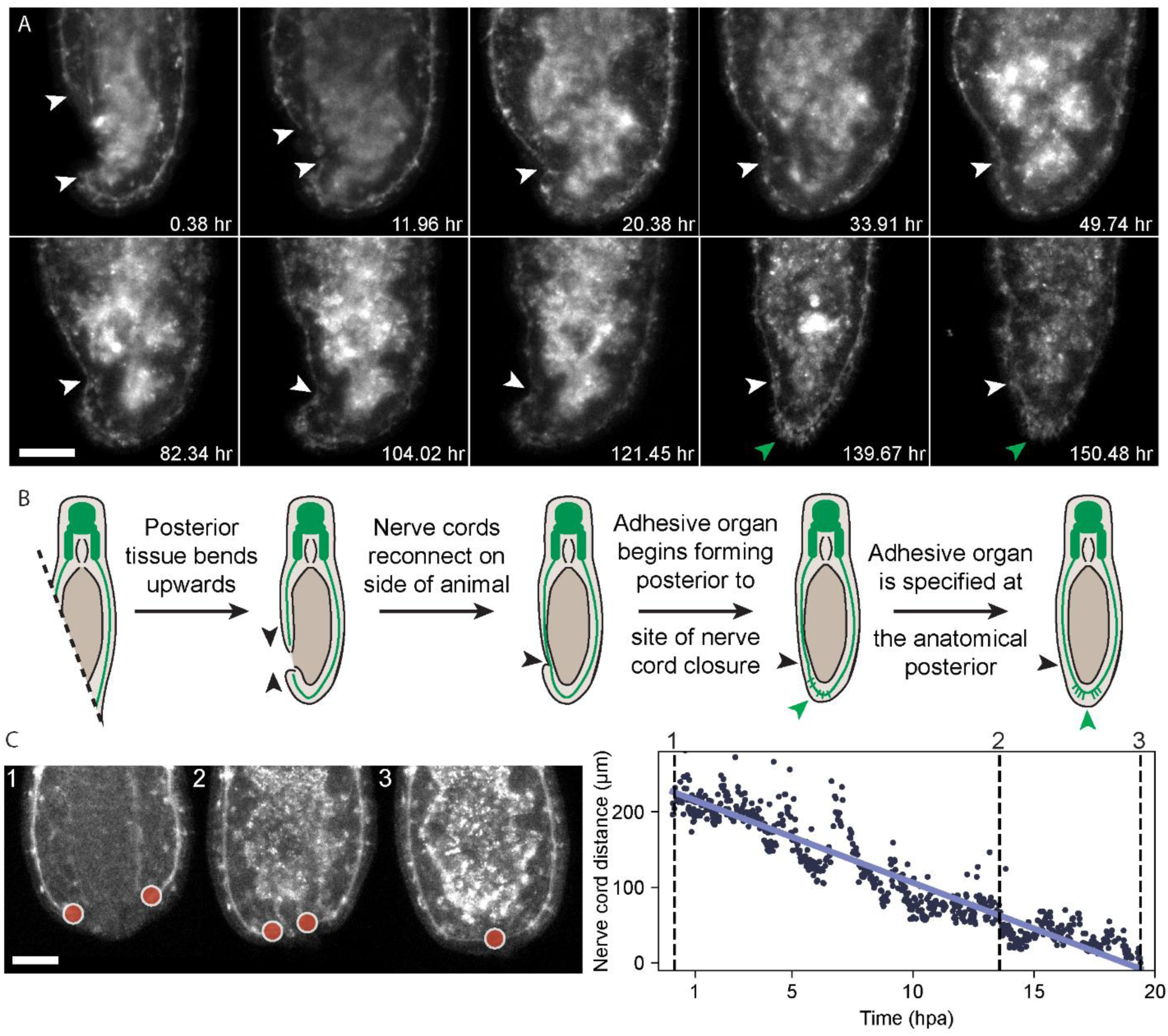
Additional characterization of nerve cord closure. Related to Figure 5 and 6. **(A)** Extreme oblique cut reveals differences in nerve cord closure and adhesive organ regeneration. Representative images of neural regeneration from a head fragment selected from a continuous week-long tracking microscopy session after an extreme (>45°) oblique amputation. White arrows: nerve cord termini and site of reconnection. Green arrow: site of adhesive organ regeneration. Scale bars: 100 µm. **(B)** Cartoon showing the process highlighting that adhesive organ regeneration occurs at a site more posterior to the point of nerve cord reconnection. **(C)** Nerve cord closure progresses at a linear rate. Nerve cord termini were manually annotated (red dots). The Euclidian distance between the two termini is plotted vs. time and exhibits a linear decrease (bottom). Dashed lines: the times corresponding to the representative images of the animal shown at the top. Scale bars: 50 µm.

**Figure S6:**
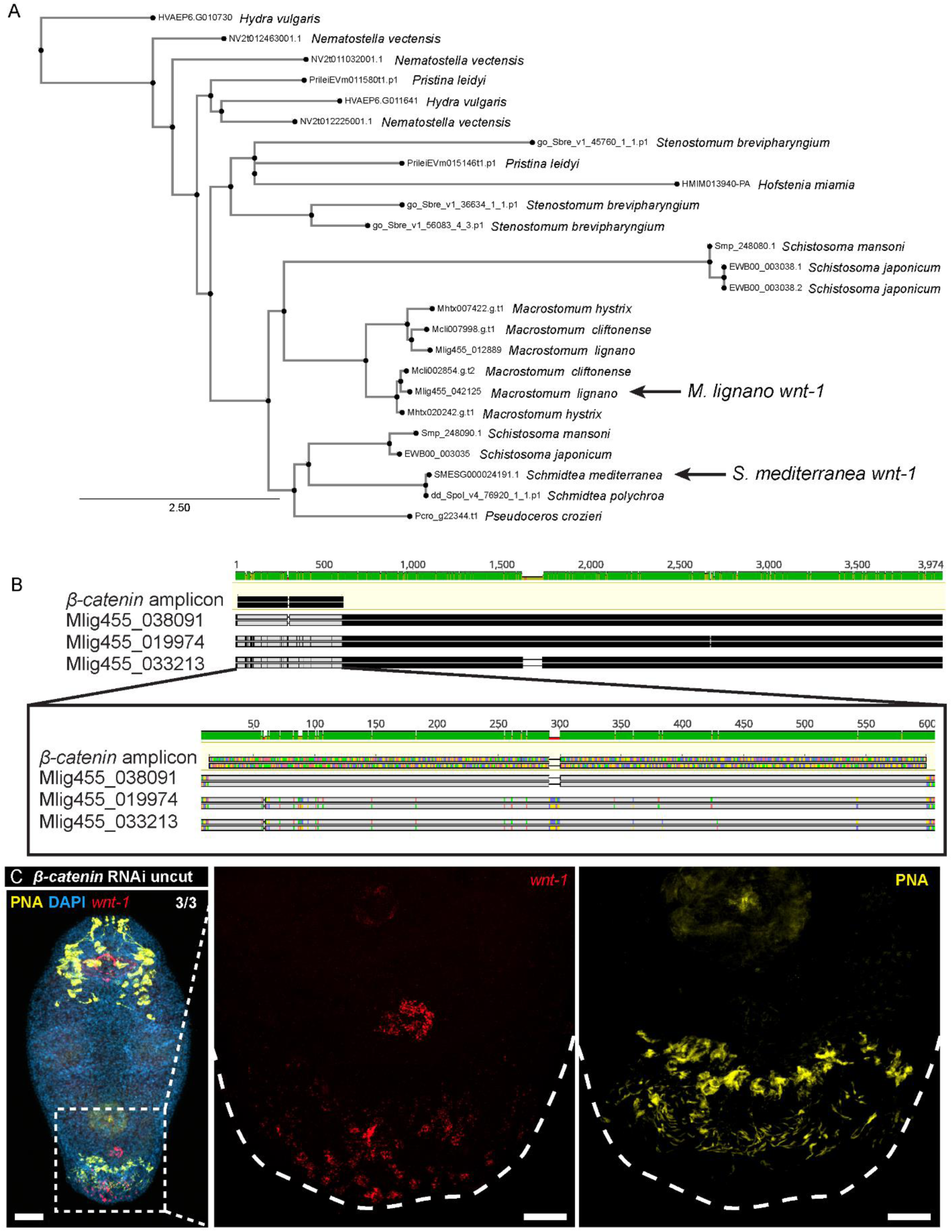
Homeostatic *wnt-1* expression remains after *β-catenin* RNAi. Related to Figure 7. **(A)** A phylogenetic tree of *wnt-1* homologs establishes *M. lignano wnt-1* as an ortholog of *S. mediterranea wnt-1.* Peptide sequences are clustered with Clustal Omega, the tree is constructed using IQTree and visualized using phylo.io. Sequences include *wnt-1* homologs in *Stenostomum brevipharyngium* (Sbre), *Schistosoma mansoni* (Smp), *Schistosoma japonicum* (EWB), *Hydra vulgaris* (HVAEP), *Nematostella vectensis* (NV), *Pseudoceros crozieri* (Pcro), and *Pristina leidyi* (Prilei), *Hofstenia miamia* (HMIM), *Macrostomum cliftonense* (Mcli), *Macrostomum hystrix* (Mhtx), *Macrostomum lignano* (Mlig), *Schmidtea mediterranea* (SMESG), *Schmidtea polychroa* (dd_Spol). **(B)** A MAFFT pairwise alignment of the region targeted by *β-catenin* RNAi to the three *β-catenin* sequences present in the genome (Mlig445_038091, Mlig445_019974, Mlig445_033213). Inset shows the broad nucleotide similarity between all three copies. Agreement with reference sequence (gray). **(C)** Homeostatic *wnt-1* expression (red) after *β-catenin* RNAi in uncut animals (left, n = 3/3). A magnified view showing *wnt-1* expression in the posterior (middle) and the adhesive glands (yellow) (right). Dashed line: outline of the tail region. Scale bars: 50 µm (left), 20 µm (middle and right).

**Figure S7:**
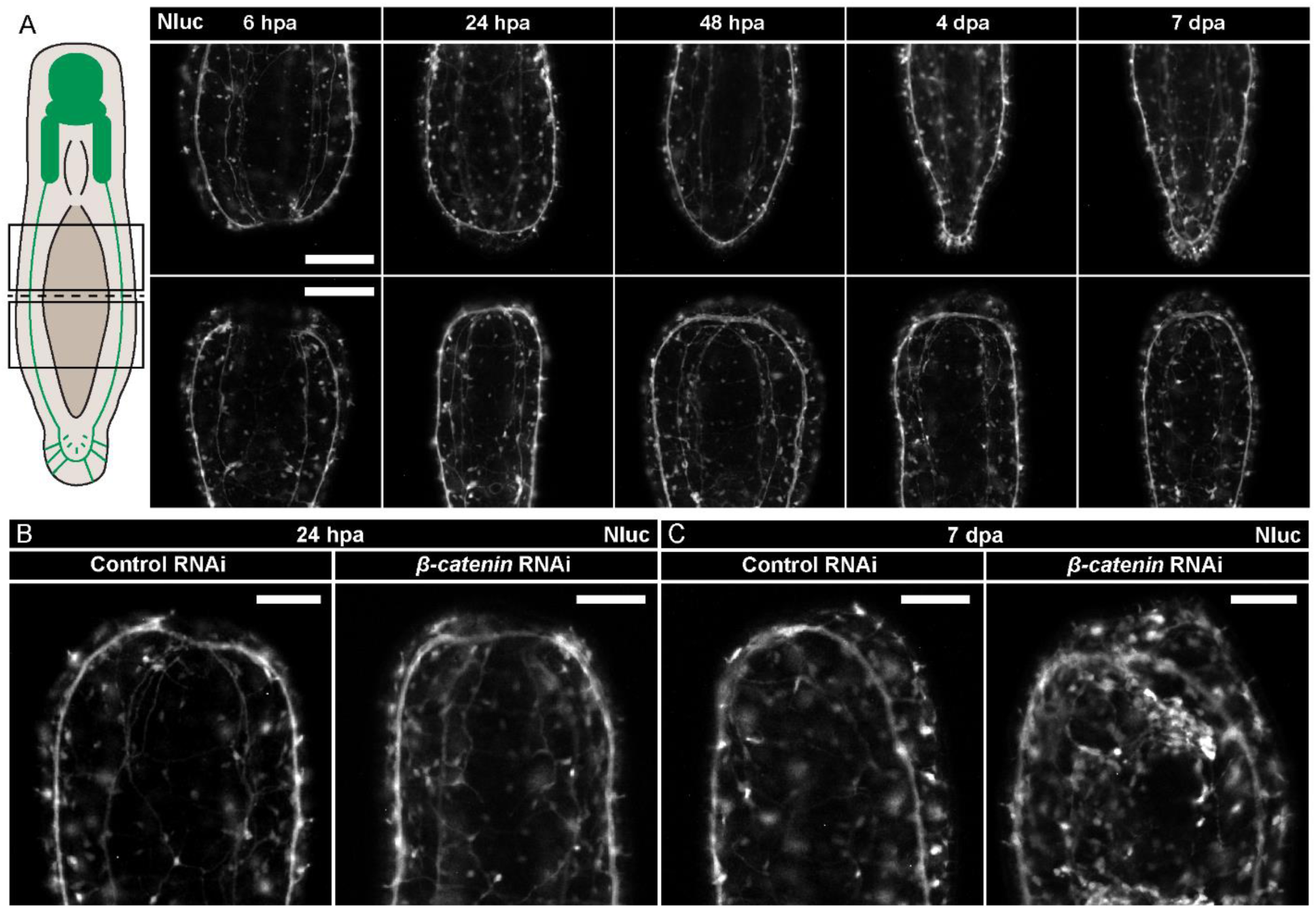
*β-catenin* RNAi fails to restore anterior regeneration. Related to Figure 7. **(A)** Representative luminescence images of neural regeneration from both head (top) and tail (bottom) fragments. Note that nerve cords are reconnected in both anterior and posterior fragments by 24 hpa. However, the posterior fragment shows no further elaboration of the neural structures even by 7 dpa. Scale bars: 50 µm. **(B-C)** Luminescence images showing anterior regeneration in control (left) and *β-catenin* RNAi (right) treated PC2 animals. At 24 hpa (D), animals in both groups heal the wound and the ventral nerve cords reconnect. At 7 dpa (E), both control animals and *β-catenin* RNAi treated animals failed to regenerate any anterior structures. Scale bars: 100 µm.

## Supplemental Tables

**Table S1: Primers, oligos, parts, and cloning utilities.** Primers for amplification of parts and TUs are provided, as well as sense and antisense primers for UNS oligo generation, the parts themselves, HCR oligo pools, and utilities for generating primers to amplify novel parts.

**Table S2: Bill of materials.** Contains key components for constructing both the luminescence/fluorescence microscope (LumiSQUID) and the tracking microscope (TrackingSQUID).

## Supplemental Videos

**Video S1:** A montage of volumetric live confocal imaging stacks showing various tissues labeled by pEnolase::GeNL.

**Video S2:** A montage of Live confocal imaging stacks showing various neural tissues labeled by pPC2::GeNL.

**Video S3:** 3D volumetric rendering of the two muscle layers wrapping the gut and body of the animal, sandwiching the ovaries.

**Video S4:** Bright field stereomicroscopy video of paralyzed pPC2::GeNL-P2A-NTR2.0::tPC2 animals treated with 5 mM MTZ for 7 days and abnormally shaped pMYH6::GeNL-P2A-NTR2.0::tSV40 animals treated with 5 mM MTZ for 7 days.

**Video S5:** Time-lapse fluorescence tracking microscope images of a two independent horizontally amputated pPC2::GeNL-P2A-NTR2.0::tPC2 animal over the course of 1 week.

**Video S6:** Time-lapse fluorescence tracking microscopy of an obliquely amputated pPC2::GeNL-P2A-NTR2.0::tPC2 animal over the course of 1 week.

## Supplemental Files

**File S1:** An archive containing plasmid maps for all parts, backbones, oligos, and additional utilities for modular assembly of transgenes.

